# CFAP410 has a bimodular architecture with a conserved surface patch on its N-terminal LRR motif for binding interaction partners

**DOI:** 10.1101/2022.09.21.508879

**Authors:** Alexander Stadler, Heloisa B. Gabriel, Laryssa V. De Liz, Santiago Alonso-Gil, Xuan Deng, Robbie Crickley, Katharina Korbula, Kaiyao Huang, Bojan Žagrović, Sue Vaughan, Jack D. Sunter, Gang Dong

## Abstract

Cilia and flagella associated protein 410 (CFAP410) is a protein localized at the basal body of cilia/flagella and plays essential roles in ciliogenesis. Multiple single amino acid mutations in CFAP410 have been identified in patients. However, the molecular mechanism for how the mutations cause these disorders remains poorly understood due to a lack of high-resolution structures of the protein. Our studies demonstrate that CFAP410 adopts a bimodular architecture. We have previously reported our structural studies on the C-terminal domain (CTD) of CFAP410 from various organisms. Here we report a 1.0-Å resolution crystal structure of the N-terminal domain (NTD) of *Trypanosoma brucei* CFAP410. We further examined how the disease-causing mutations in this domain may affect the folding and structural stability of CFAP410. Our results suggest that the single-residue mutations in the CFAP410-NTD cause human diseases by destabilizing the structure that subsequently disrupts its interaction with other partners.

## Introduction

Cilia- and flagella-associated protein 410 (CFAP410) was originally named C21orf2 as it is coded by a gene that was mapped to chromosome 21 open reading frame 2 [1]. Immunofluorescence analysis showed that CFAP410 localizes to the basal body and works together with NEK1 and SPATA7 as a functional module in ciliogenesis and DNA damage repair [2–4]. CFAP410 exhibits significant sequence conservation across phyla and is present in all sequenced genomes of ciliates. *Homo sapiens* CFAP410 (HsCFAP410) consists of 255 amino acids. A number of single residue mutations of HsCFAP410 have been identified in patients with skeletal and/or retinal disorders, including I35F, C61Y, R73P, Y107C, Y107H, V111M and L224P [2, 5–8]. These disorders are genetically autosomal recessive and manifested as dysplasia of the ribs, vertebral bodies, ilia, proximal femora in skeletons, and retinitis pigmentosa in eyes [9, 10]. However, it remains unknown how these single amino acid mutations in HsCFAP410 cause so deleterious defects.

We have carried out extensive structural and biochemical characterizations on three homologs of CFAP410 from *H. sapiens, Trypanosoma brucei* and *Chlamydomonas reinhardtii.* Our work revealed that CFAP410 is a bimodular protein comprising two highly conserved domains, including a leucine rich repeat (LRR) motif at its N-terminus and an oligomeric helical bundle at its C-terminus. We have recently reported crystal structures for the C-terminal domain (CTD) of the three homologs of CFAP410 and how the disease-causing L224P mutation abolishes its localization to the basal body (*Open Biol.* in press). Here we report our further structural studies on the N-terminal domain (NTD) of *T. brucei* CFAP410 (TbCFAP410). The structure allows us to examine how the mutations might affect the folding of the protein. We further checked the localization of the wild-type and disease-causing mutations in *T. brucei* and analyzed its potential interaction with NEK1 via a conserved surface patch. Taken together, our work provides an explanation how single-residue mutations in CFAP410 might cause ciliopathies.

## Results

### Crystal structure of TbCFAP410-NTD reveals a canonical LRR motif with a highly conserved surface patch

Our bioinformatic analysis suggests that all CFAP410 orthologs have a bimodular organization consisting of two folded domains that are connected by a disordered loop. We have previously determined high-resolution crystal structures for the CTD of three CFAP410 proteins from *H. sapiens*, *T. brucei* and *C. reinhardtii* (*Open Biol.* in press). Here, we report our further investigation on the NTD of CFAP410. We made multiple truncation constructs based on our bioinformatic analysis and tried different fusion tags, including His6, SUMO, and maltose-binding protein (MBP). No soluble proteins could be obtained for the constructs of HsCFAP410 and CrCFAP410. For TbCFAP410 we managed to produce soluble proteins for four of the six truncations we generated, which cover residues 1-177, 1-170, 1-164 and 1-160, whereas two other shorter constructs spanning residues 1-149 and 1-140 were insoluble upon overexpression in bacteria.

Purified proteins of all these four soluble TbCFAP410-NTD truncations were subjected to extensive crystallization screenings. In the end we obtained crystals only for the construct spanning residues 1-160, which is what NTD refers to in all subsequent structural analyses of TbCFAP410. Single crystals of TbCFAP410-NTD diffracted X-rays to approximately 1.0 Å resolution at the ESRF synchrotron site (**Table 1**). The crystal structure was determined by the molecular replacement method using a homology model of residues 16-138 of TbCFAP410 generated by SWISS-MODEL [11] based on the crystal structure of the *C. reinhardtii* axonemal dynein light chain-1 (PDB code: 6L4P) [12].

**Table 1.** Data collection and refinement statistics.

|  |  |
| --- | --- |
| Beamline | ESRF ID23-1 |
| Wavelength (Å) | 0.972 |
| Resolution range (Å) | 19.90 - 1.0<br>(1.04 - 1.0) |
| Space group | P 2<sub>1</sub> 2<sub>1</sub> 2<sub>1</sub> |
| Unit cell<br>*a*, *b*, *c* (Å)<br>$\alpha$, $\beta$, $\gamma$ (°) | 39.80, 52.96, 61.43<br>90, 90, 90 |
| Total reflections | 789,670 (30,477) |
| Unique reflections | 69,533 (5,835) |
| Multiplicity | 11.4 (5.2) |
| Completeness (%) | 98.17 (83.75) |
| Mean *I*/$\sigma$(*I*) | 10.84 (0.61) |
| R-merge | 0.099 (1.695) |
| R-meas | 0.104 (1.882) |
| R-pim | 0.030 (0.792) |
| CC<sub>1/2</sub> | 0.998 (0.386) |
| CC* | 1.00 (0.746) |
| Reflections used in refinement | 69,519 (5,834) |
| Reflections used for R-free | 1,989 (168) |
| R-work | 0.152 |
| R-free | 0.162 |
| Number of heavy atoms in proteins | 1,302 |
| Number of solvent | 157 |
| RMS (bonds) | 0.010 |
| RMS (angles) | 1.22 |
| Ramachandran favored (%) | 96.23 |
| allowed (%) | 3.77 |
| outliers (%) | 0.00 |

Statistics for the highest-resolution shell are shown in parentheses.

The refined model shows that all residues in the construct could be traced and confidently built (**Fig. 1A**). Residues 1-138 fold into a canonical leucine-rich repeat (LRR), with five tandem repeats forming a calzone-like structure. The LRR motif contains five parallelly arranged β strands forming a flat sheet on one side and an array of four short α helices on the other. The space between the α helices and the β strands constitutes the hydrophobic core of the LRR motif and is packed with many highly conserved leucine residues. Additionally, there is a short α helix on either end of the LRR motif that acts as a cap and is oriented roughly perpentidular to the β sheet. The overall structure is slightly bent toward the β sheet side as seen in other LRR structures [13]. Additionally, we found that residues 139-160 of TbCFAP410 forms a long hairpin-like loop, which is tightly packed on top of the helical surface of the LRR motif (**Fig. 1B**). The loop makes extensive contacts with the LRR motif via six hydrogen bonds and numerous hydrophobic interactions (**Fig. S1A**). Those residues in the loop involved in its interaction with the LRR motif are highly conserved in all homologs from various kinetoplastids, so as the interacting residues on the LRR side (**Fig. S1B-D**).

**Figure 1.**
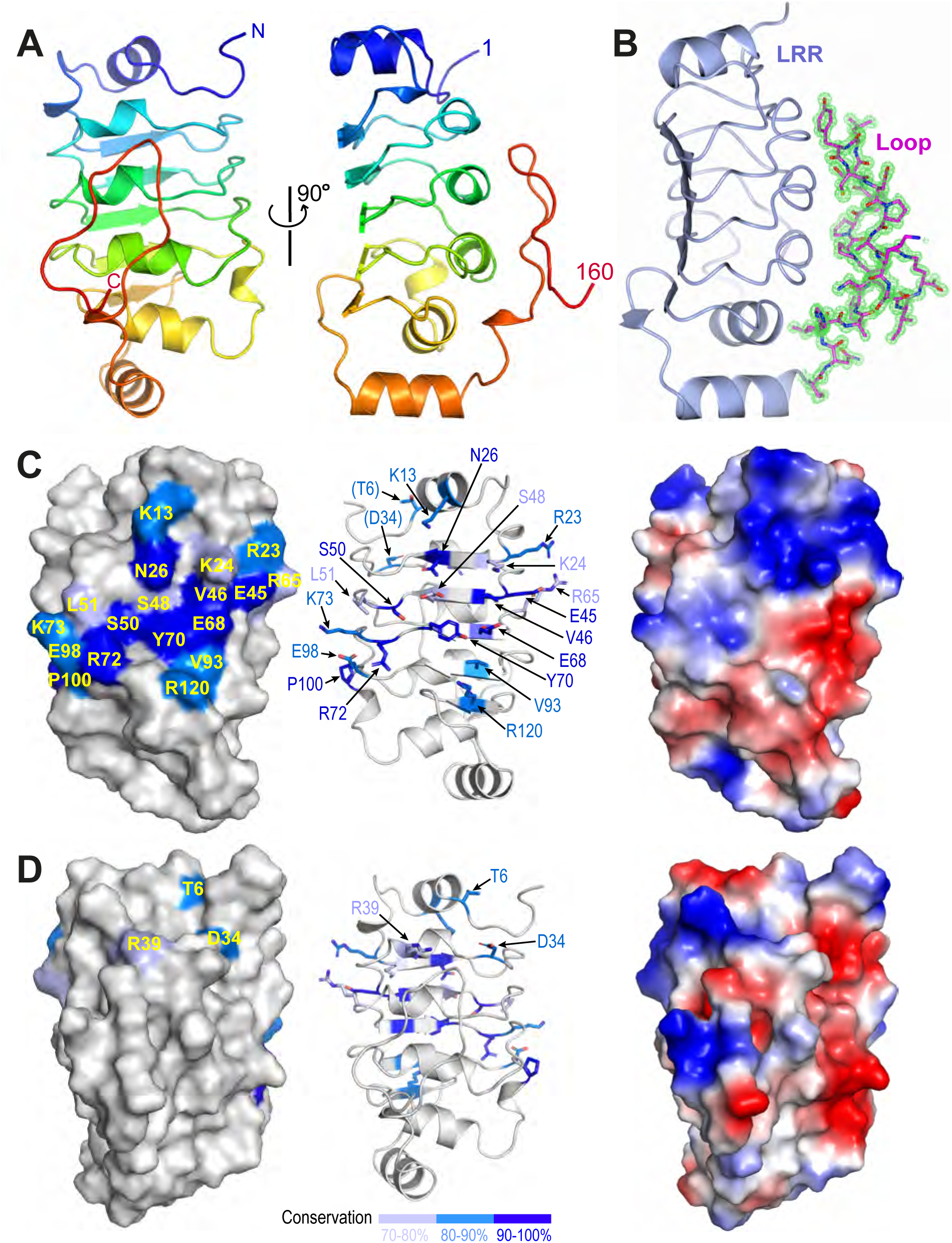
Crystal structure of the N-terminal LRR motif of TbCFAP410. (**A**) Ribbon diagram of TbCFAP410-NTD (aa1-160) color-ramped from blue (N-terminus) to red (C-terminus). (**B**) Cartoon of TbCFAP410-NTD with its C-terminal loop (aa141-160) shown as sticks together with 2*F_o_*-*F_c_*electron density map contoured at 1.5 σ level. (**C**) A conserved surface patch on the flat β sheet of the LRR motif. Conserved surface residues are colored in blue, marine and light blue based on their sequence identities. (**D**) The opposite side of the LRR motif covered by the loop is not conserved. An electrostatic surface plot in the same orientation is shown at the right side in both (C) and (D).

Sequence alignments of CFAP410 homologs from mammals, trypanosomes and algae reveal an extensive surface patch on the flat β sheet of the LRR motif. The patch includes eighteen residues that are highly conserved in all homologs across these phyla (**Fig. 1C**; **Fig. S2A, B**). The opposite side of the LRR motif is covered by a long hairpin-like loop in TbCFAP410 as shown above. Although this loop is conserved in kinetoplastids, it is essentially devoid of any conserved residues in all other homologs from animals and algae (**Fig. 1D**; **Fig. S2C**).

### Examination of disease-causing mutations on CFAP410-NTD

Multiple single-residue mutations in HsCFAP410 have been identified in patients. Except for L224P located at the CTD, all other mutations are in the N-terminal LRR motif of HsCFAP410, including I35F, C61Y, R73P, Y107C, Y107H and V111M [2, 5–8]. Sequence alignment shows that the LRR motif is highly conserved in animals, trypanosomes and algae (**Fig. S2A**). Except for R73, all sites associated with disease-causing mutations are highly conserved hydrophobic residues. The LRR motifs of *H. sapiens* and *T. brucei* CFAP410 share 41% identity and 60% similarity (**Fig. S3A**). Comparison of the crystal structure of TbCFAP410-NTD with structural models of HsCFAP410 (aa1-130) generated by SWISS-MODEL (template-based) [11] and AlphaFold2 (*ab initio*) [14] shows that they all share a highly similar three-dimensional structure (**Fig. S3B**). Particularly, all the five disease-causing mutation sites mentioned above adopt essentially the same conformation in these structures. Statistics reports show that both predicted models of HsCFAP410 were built with high confidence (**Fig. S3C-E**). These allow us to examine how the mutations of single residues in HsCFAP410 might affect its structure and function based on the crystal structure we have determined for TbCFAP410-NTD.

Location of individual residues corresponding to the five disease-causing mutations in HsCFAP410 can be clearly visualized in TbCFAP410 (**Fig. 2A, B**). These residues can be grouped into three categories, including one surface residue that is fully exposed to solvent (R75), two semi-exposed residues (V37 and Y107), and two hidden residues that are completely buried in the hydrophobic core of the structure (C63 and V111) (**Fig. 2C-E**).

**Figure 2.**
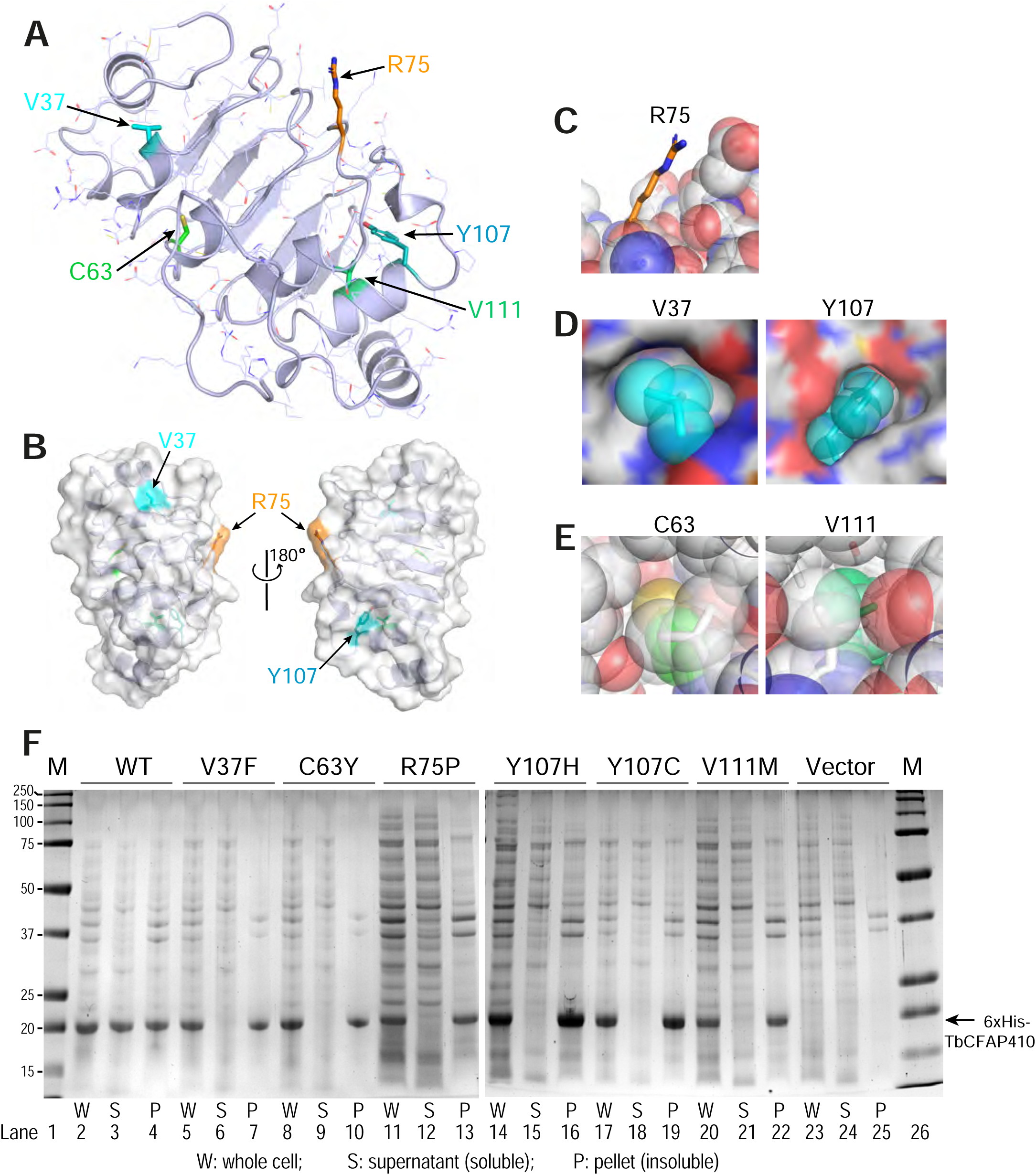
Structural environment and expression test of the disease-causing mutations in TbCFAP410-NTD. (**A**) Crystal structure of TbCFAP410-NTD with residues corresponding to disease-causing mutations in HsCFAP410 shown as sticks and labeled. All other side-chains are shown as lines. (**B**) Front and back views of TbCFAP410-NTD shown as semi-transparent surface and ribbons. Only three of the five residues in (A) exposed to solvent are labeled. (**C**-**E**) The five disease-causing mutation sites are grouped into three categories: fully exposed (C), semi-buried (D), and fully buried (E). (**F**) Solubility test of wild-type and mutants of TbCFAP410-NTD upon over-expression in *E. coli*. Expression with only the vector is included as a negative control.

Given that most of these disease-causing mutants are hydrophobic and partially or fully buried in the structure, we therefore examined whether these mutations affect the folding or structural stability of the protein. Wild-type and mutant TbCFAP410-NTD proteins with an N-terminal His6 tag were expressed in *Escherichia coli*. After breaking the cells and spinning down the cell lysate, we checked the soluble fractions in the supernatant and the insoluble parts in the pellet. Our data show that wild-type TbCFAP410-NTD were present in both fractions, suggesting that it is at least partially soluble (**Fig. 2F**, lanes 2-4). In contrast, however, all the six mutants were present only in the pellets but not in the supernants (**Fig. 2F**, lanes 5-22), demonstrating that the proteins are insoluble and thus likely mis-folded.

We further examined the crystal structure trying to understand how these mutations may cause changes in the folding of the protein. Firstly, mutation R73P of HsCFAP410, equivalent to residue R75 in TbCFAP410, is located in a short linker tightly packed between an α helix and a β strand in the structure (**Fig. 3A**). We found that its fully solvent exposed side-chain forms a stacking interaction with that of residue E53 via two salt bridges. Additionally, the amide proton of residue R75 forms a hydrogen bond with the carbonyl group of residue L54. Mutating R75 to proline would disrupt all these interactions. Furthermore, the circular side-chain of proline in the mutant would additionally cause clashes with the neighboring residues in the tightly packed region of the structure, and thus destabilizes the structure of the protein.

**Figure 3.**
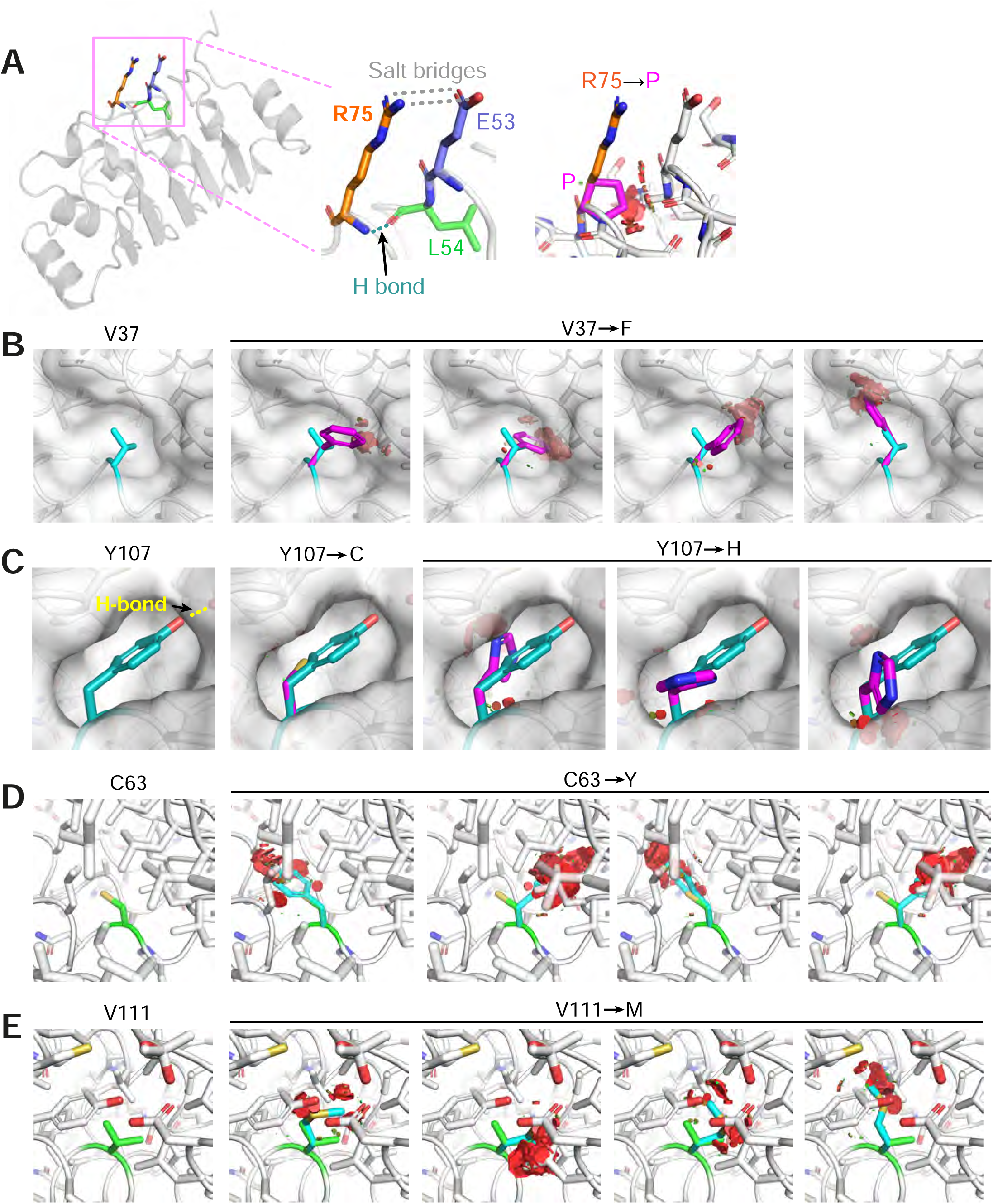
Structural examination of how the disease-causing mutants on TbCFAP410-NTD may affect protein folding or structural stability. (**A**) Residue R75 is in a rigid linker and forms hydrogen bonds with residues from the neighboring linker. Its mutation to proline would disrupt the salt bridges and the hydrogen bond, as well as cause severe clashes with neighboring residues. (**B**) Residue V37 is located in a small cleft. Its replacement by phenylalanine would clash with the surrounding residues. (**C**) Residue Y107 locates in a semi-open eggshell-like pocket and the hydroxyl group on its side-chain forms a hydrogen bond with a neighboring residue. Both Y107C and Y107H mutants would disrupt this hydrogen bond and generate a cavity or clash with other residues surrounding the pocket. (**D**, **E**) Residues C63 and V111 are both buried deeply in the hydrophobic core of the structure. Their replacements by residues with bulkier side-chains (i.e. tyrosine and methionine) would cause sever clashes with the neighboring residues. All images were generated using Pymol (http://www.pymol.org).

All the other five ciliopathy mutation sites within HsCFAP410-NTD occur at highly conserved hydrophobic residues (**Fig. S2A**). Our structural analyses on the equivalent residues in TbCFAP410-NTD show that residue V37 (corresponding to I35 in HsCFAP410) is packed tightly against a small hydrophobic cleft. Its replacement by phenylalanine (i.e. I35F in HsCFAP410) would cause multiple clashes between the bulky aromatic side-chain and the neighboring residues around the cleft (**Fig. 3B**). Regarding the two mutations Y107C and Y107H, residue Y107 is located in a semi-open eggshell-like pocket. The hydroxyl group of Y107 forms a hydrogen bond with the side-chain of a neighboring residue. Mutation Y107C would apparently disrupt this hydrogen bond as well as generate a void cavity (**Fig. 3C**). It is similar for mutation Y107H, but this mutation would additionally cause clashes between the side-chain of His and other hydrophobic residues around the pocket. For the other two mutants C63Y and V111M, both residues are fully buried inside the hydrophobic core of the structure (**Fig. 3D, E**). Their replacements by residues with a bulkier side-chain would inevitably cause severe clashes and thus disrupt the folding of CFAP410-NTD. Our molecular dynamics simulation analyses indeed demonstrated significant changes in all these disease-causing mutants, particularly C63Y, in comparison with the crystal structure of wild-type TbCFAP410-NTD (**Fig. S4A, B**). Strutural modeling by AlphaFold2 suggests that mutant C63Y would cause the flip of an α helix in the LRR motif to accommodate the large aromatic sidechain of tyrosine [14]. Such flip would subsequently distablizes the interaction between the helix and the hairpin loop to cause the exposure of the hydrophobic residues covered by the loop (**Fig. S4C**).

### *In vivo* localization of the disease-causing mutants of TbCFAP410

To examine the impact of these disease-causing mutations on the function of TbCFAP410 *in vivo,* we generated individual mutant constructs with an N-terminal mNeonGreen fluorescent protein tag and expressed them in *T. brucei* using a tetracycline inducible expression system. All expressions were induced for 24 h. The cells, which also expressed SAS6 endogenously tagged with mScarlet as a marker of the basal body and pro-basal body, were then examined by wide-field fluorescence microscopy (PMID: 33165561). The wild-type protein localized to the basal and pro-basal bodies as well as the posterior cell tip, together with a weak cytoplasmic signal (**Fig. 4A**, left), matching the localization observed in the TrypTag project [15]. We found that the localization of the mutants in the NTD was similar to that of the wild-type CFAP410, with a posterior cell tip signal and a basal body signal but a variable signal from the cytoplasm. Additionally, we examined a single central amino acid mutation (Y70A) and two clustered mutations within the surface patch that is highly conserved across all phyla, either mutating only the central ridge of the patch (4 mut: E68K/Y70A/R72E/K73E) or a more extensive one with two extra flanking residues mutated as well (6 mut: E68K/Y70A/R72E/K73E/N26A/R120E). The localization patterns of these mutants were also similar to that of the wild-type protein (**Fig. 4A**, left).

**Figure 4.**
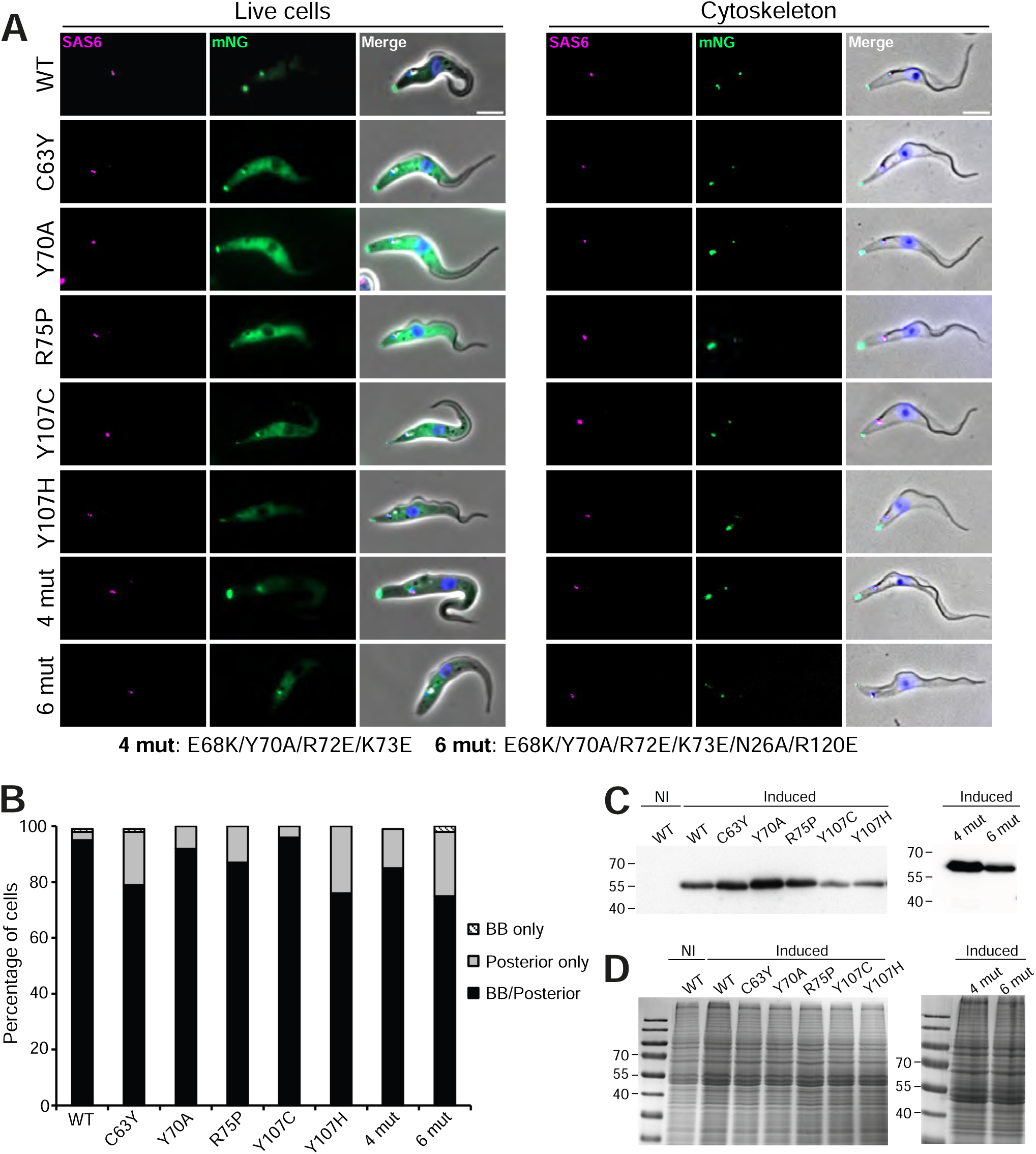
Fluorescence microscopy images from wild-type and mutants of TbCFAP410. (**A**) Images acquired from live cells or methanol fixed cytoskeletons from doxycycline induced cells expressing TbCFAP410 wild-type or TbCFAP410 mutants (green) with mSc::TbSAS6 - basal body marker (magenta). Scale bar: 5 µm. (**B**) Graph of TbCFAP410 signal counts. The percentage of cytoskeletons with a basal body and posterior signal, a basal body only signal and posterior signal only was calculated for each cell line. Induction was repeated twice, with a similar result both times and data from one is shown. BB: basal body. N=100 cells. (**C**) Western blot showing the induction of expression of wild-type and mutant TbCFAP410 in *T. brucei*. BB2 (anti-Ty) antibody was used. 4×10^6^ parasites per lane were loaded. NI: non-induced RNAi. (**D**) Coomassie stained polyacrylamide gel of samples loaded in (**C**).

To further investigate how stably the mutated proteins were associated with the cytoskeleton, we next generated detergent extracted cytoskeletons of the induced cells and quantified the localization of these mutants. Similar to what was seen in live cells, the wild-type and mutant protein generally remained stably associated with both the basal body and the posterior cell tip (**Fig. 4A**, right). The distribution of the localization of wild-type and all mutant TbCFAP410 was quantified (**Fig. 4B**). For all mutants except Y107C, there was a slight reduction in the localization of the protein to the basal body. Expression of the wild-type and all mutant proteins after induction was confirmed by western blot (**Fig. 4C, D**).

### Depletion of CFAP410 causes no significant changes in ciliogenesis in *C. reinhardtii*

To check if CFAP410 plays an essential role in ciliary assembly in another organism, we carried out depletion tests in *C. reinhardtii*. We successfully generated two targeted knockout mutants of CrCFAP410 using CRISPR/Cas9 (**Fig. 5A-C**). However, although there was notable reduction in both flagellar length and percentage of flagellated cells in the two knockout cell lines in comparison to the wild-type cells, the differences were not as dramatic as what would be expected for an essential protein (**Fig. 5D, E**). Interestingly, we found there is another much shorter isoform of CrCFAP410, which we named “CrCFAP410-short”. CrCFAP410-short contains exactly the same N-terminal region (aa1-132) as CrCFAP410, but it lacks the connecting loop and the whole C-terminal oligomeric domain present in CrCFAP410 (**Fig. 5F**). We hypothesize that CrCFAP410-short may work redundantly to CrCFAP410 and could thus compensate the depleted CrCFAP410 to maintain normal flagellar structure and function.

**Figure 5.**
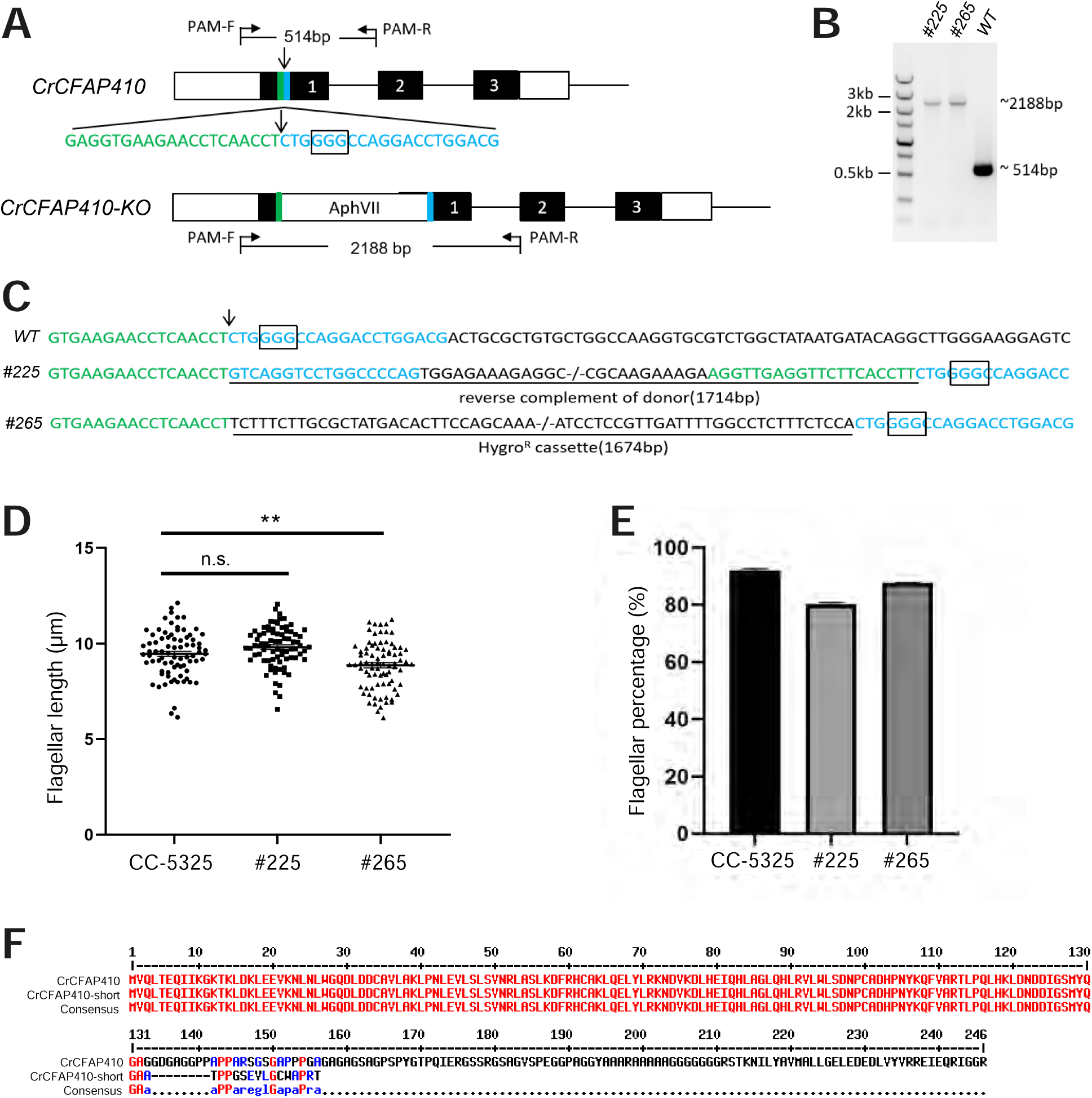
Targeted knockout of CrCFAP410 by co-delivery of RNP complex and microhomologous dsDNA donor. (**A**) Schematic of the CrCFAP410 loci with dsDNA donor carrying a hygromycin resistance cassette inserted into the Cas9 cleavage site. Exons are indicated as black boxes, together with 5′ and 3′ UTR (white boxes), the cleavage site (arrow), the PAM site (the rectangular box) and 20 bp homologous arms (green and blue). The predicted sizes of the PCR fragments were 514 bp and 2,188 bp using PCR primers PAM-F and PAM-R in wild-type CC-5325 and in the knock-out mutant, respectively. (**B, C**) PCR results (B) and DNA sequences of PCR products (C) for wild-type and two mutants (#225 and #265). The 1,714 bp reverse complement of donor containing the two homologous arms was imprecisely integrated into the Cas9 cleavage site in mutant #225. The 1,674 bp hygromycin resistance cassette was precisely inserted in mutant #265. The insertion at the cleavage site was underlined. (**D, E**) The percentage of flagellated cells (D) and length (E) in CC-5325, *CrCFAP410* knock-out mutants #225 and #265. Data are represented as mean ± standard error (S.E.). Statistical analysis was performed using a two-tailed unpaired Student’s *t*-test. **p<0.005, n.s., not significant. (**F**) Sequence alignment of CrCFAP410 and CrCFAP410-short.

## Discussion

CFAP410 is a protein present in all sequenced ciliates and plays an essential role in ciliogenesis [2]. Located mainly at the basal body, the protein is highly conserved across phyla including algae, protists and animals. Our analysis shows that CFAP410 is a bimodular protein comprising two folded domains, i.e. NTD and CTD, which are connected via a variable unstructured loop. All single-residue mutations causing ciliopathies are located in the NTD or CTD. We have previously reported crystal structures of the CTD of three CFAP410 homologs from *T. brucei*, *C. reinhardtii*, and *H. sapiens*, which all adopt a similar tetrameric conformation (*Open Biol.*, in press). Multiple single-residue mutations in HsCFAP410 have been identified in patients with SMS or RDMS diseases [6–8]. Except for L224P, all other six ciliopathy mutations locate in the NTD of HsCFAP410. Here we report a high-resolution crystal structure of TbCFAP410-NTD, which shares 41% identities with the HsCFAP410. Comparison of our crystal structure with the predicted models of HsCFAP410 demonstrates a similar conformation. Therefore, we could extrapolate from the structure of TbCFAP410-NTD how the disease-causing mutations might affect the structure and function of HsCFAP410.

To check how the disease-causing mutations affect the function of CFAP410 in the native environment, we further carried out *in vivo* assays on TbCFAP410. Our results showed that unlike the mutant L272P that completely lost its localization at the basal body and the posterior cell tip, none of the mutants in the NTD showed significant change in their localization in the cell. Our solubility tests for recombinant protein in *E. coli* demonstrate that all the mutants in the NTD of TbCFAP410 cause structural instability as mutant proteins were completely insoluble upon overexpression in *E. coli*. However, the defect in the folding of the NTD did not affect the specific localization of TbCFAP410 in the cell. Moreover, we did not observe stable interaction between the individually purified NTD and CTD of TbFAP410 (data not shown). All these strongly suggest that the two modules in CFAP410 likely function independently, with the CTD controlling its localization and the NTD regulating its other functions.

In the crystal structure of TbCFAP410-NTD there is an extensive surface patch across all five strands of the β sheet. This patch consists of eighteen highly conserved residues. We speculate that this surface patch might play an important role for the function of CFAP410. However, none of the mutations within the patch we tested showed significant defects in cellular localization of the protein in *T. brucei*. We think that the function of the conserved patch in CFAP410 is for its other functions. Previous studies have shown that CFAP410 is a component of a retinal ciliopathy-associated protein complex containing both NEK1 and SPATA7 [16]. Direct interaction between CFAP410 and the C-terminal end of NEK1 has been observed in human cells [17]. We therefore speculate that the conserved surface patch on the NTD of CFAP410 is possibly involved in its interaction with NEK1. We carried out structural modelling using AlphaFold-multimer for the complex of HsCFAP410-NTD (aa1-150) and HsNEK1-CTD (aa1208-1286). In the prediction, the multiple sequence alignment (MSA) identified approximately 3,000 and 1,000 sequences similar to HsCFAP410-NTD and HsNEK1-CTD, respectively. The five generated models of the complex could be perfectly superimposed (**Fig. 6A**). The predicted local distance difference test (pLDDT) score was above 90 for most parts of both molecules, indicating very high accuracy that is equivalent to structures determined by experiments (**Fig. 6B**). The prediction aligned error (PAE) score, which displays the calculated errors for the predicted distance of each pair of residues, was also substantially low for the residues on the interface in all the five models (**Fig. 6C**). All these suggest that the conserved surface on the NTD of CFAP410 is likely the binding site of NEK1.

**Figure 6.**
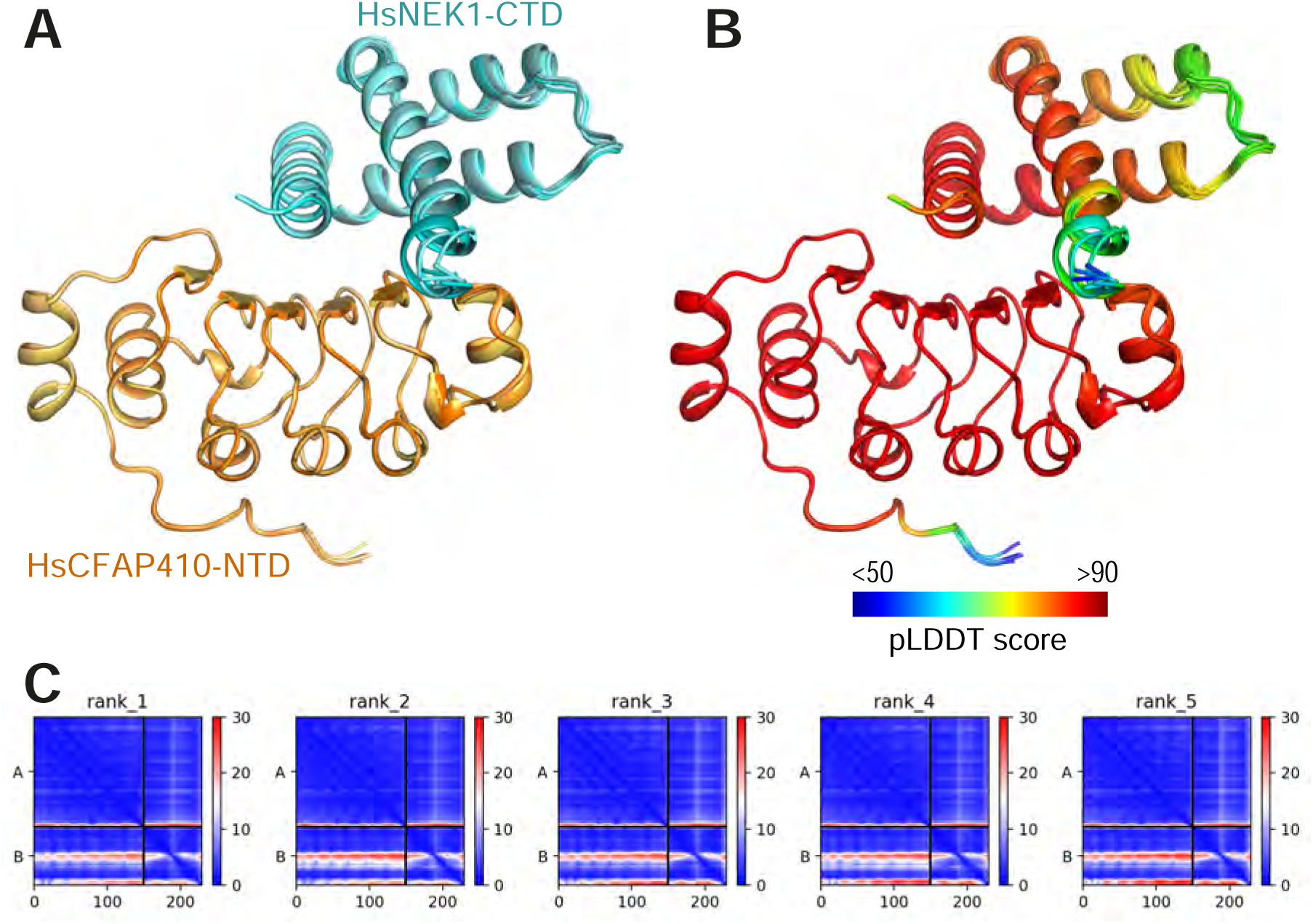
Predicted interaction between HsCFAP410-NTD and HsNEK1-CTD. (**A**) Superimposed cartoons of the top-ranked five models of the HsCFAP410-NTD (aa1-150) and HsNEK1-CTD (aa1208-1286) complex generated by AlphaFold2-multimer. (**B**) The same view as in (A) but colored by the predicted local distance difference test (pLDDT) score at each position. (**C**) Prediction aligned error (PAE) score for the five models, which indicates the calculated error of the predicted distance for each pair of residues. The uncertainty in the predicted distance of two amino acids is color coded from blue (0 Å) to red (30 Å), as shown in the right bar. The upper left quadrant corresponds to errors in the distances of residues within HsCFAP410-NTD, the lower right quadrant to errors within HsNEK1-CTD, and both the upper right and lower left quadrants to errors between CFAP410 and NEK1. The differences among the five models are rather marginal.

Although the helical side of the LRR motif lacks conservation across phyla, a cross-species conservation pattern exists in all kinetoplastids. The hairpin-like loop in the crystal structure of TbCFAP410-NTD spans over 20 residues (aa139-160) and makes extensive contacts with the LRR motif via multiple hydrogen bonds as well as numerous hydrophobic interactions. Those residues in the loop involved in its interaction with the LRR motif are highly conserved in all kinetoplastid homologs, so as the corresponding interaction residues on the LRR motif. All these suggest that this extensive loop is a unique feature present only in kinetoplastid species. As mentioned above, all TbCFAP410 truncations with the intact loop were soluble, whereas those lacking parts or the whole loop, i.e. constructs covering residues 1-149 and 1-140, were all insoluble upon expression in *E. coli*. This further suggests that the loop is essential to facilitate the folding and/or to stablize the structure of the LRR motif in TbCFAP410. The stabilizing loop might be an ancient element retained during evolution in the early branched flagellated Kinetoplastida.

CFAP410 is a ciliary protein localized to the base of cilia in human cells and involved in ciliopathies characterized with retinal and skeletal impairment [2, 7]. Previous comparative proteomics studies show that the protein level of CFAP410 increased by 1.59 fold in disassembling flagella of *C. reinhardtii* [18]. We therefore investigated whether CFAP410 plays an essential role in ciliogenesis. We generated two targeted knockout mutants of CFAP410 in *C. reinhardtii*. However, we did not observe dramatic change in either flagellar length or percentage of flagellated cells in comparison to the wild-type cells.

This might be explained by the presence of another short version of CrCFAP410 that lacks the whole C-terminal oligomeric domain. The latter may work redundantly to CrCFAP410 and thus compensate the deletion of CrCFAP410.

Similar to *C. reinhardtii*, there is also a paralog of CFAP410 in *T. brucei* (Tb927.10.5130). This paralog also localizes to the basal body, pro-basal body and posterior cell tip in same way as TbCFAP410 [15]. Sequence alignment shows that the paralog has a similar size to TbCFAP410 and their N-terminal regions adopt a similar conformation (**Fig. S5A, B**). Furthermore, we found that most residues within the surface patch on TbCFAP410 are also conserved in the paralog (**Fig. S5C, D**). It is possible that the two proteins might function redundantly in *T. brucei*.

Although only a single paralog or truncated form of CFAP410 is present in *T. brucei* and *C. reinhardtii*, there are in total eleven isoforms of CFAP410 in human cells. In contrast to the two related proteins in *T. brucei* and *C. reinhardtii* that either misses the CTD or contains a very different CTD from that present in CFAP410, all the eleven HsCFAP410 isoforms contain an identical CTD (**Fig. S6A**). But these isoforms have diversified in their N-terminal part or the linker region that connects the two folded domains. A tetrameric model of full-length HsCFAP410 generated by AlphaFold-multimer showed that the CTD forms the same tightly packed helical bundle as revealed by the crystal structures, with the N-terminal LRR motifs of the four molecules loosely linked the bundle via a long unstructured loop (**Fig. S6B**). The large number of CFAP410 isoforms in humans might be formed from alternative splicings, variable promoter usage, or other post-transcriptional modifications of the same gene, with each unique mRNA sequence subsequently produces a specific form of the protein [19–21]. Whether these isoforms are expressed in a tissue-specific manner like many other human proteins needs to be further investigated in the future. The fact that they all contain the same CTD that forms an oligomeric structure suggests that this domain likely plays an important role for the function of HsCFAP410 in humans. Despite variability in the the connecting loops, most of these isoforms also contain an intact NTD with nearly identical sequences in their structural cores. It further suggests that the NTD likely also plays a critical role in their function.

In summary, taken together our extensive structural, biochemical and *in vivo* studies, we conclude that CFAP410 forms a tetramer via its CTD that forms a tightly packed eight-helix bundle, which gathers the four globular NTDs together to form a clustered protein complex (**Fig. 7**). The NTD of CFAP410 adopts the conformation of a canonical LRR motif and contains an extended surface patch. The patch is highly conserved across distant phyla including algae, protist parasites and mammals, and thus likely plays an essential role in the *in vivo* function of CFAP410, such as binding to NEK1 or other interaction partners. The molecular mechnism for how the single-residue mutations in HsCFAP410 cause human diseases seems to be either the disruption of the oligomeric assembly of the CTD (for mutation L224P), which abolished its correction localization in the cell, or destabilizing the structure of the NTD (for mutations I35F, C61Y, R73P, Y107C, Y107H and V111M) that disrupts its interaction with other partners.

**Figure 7.**
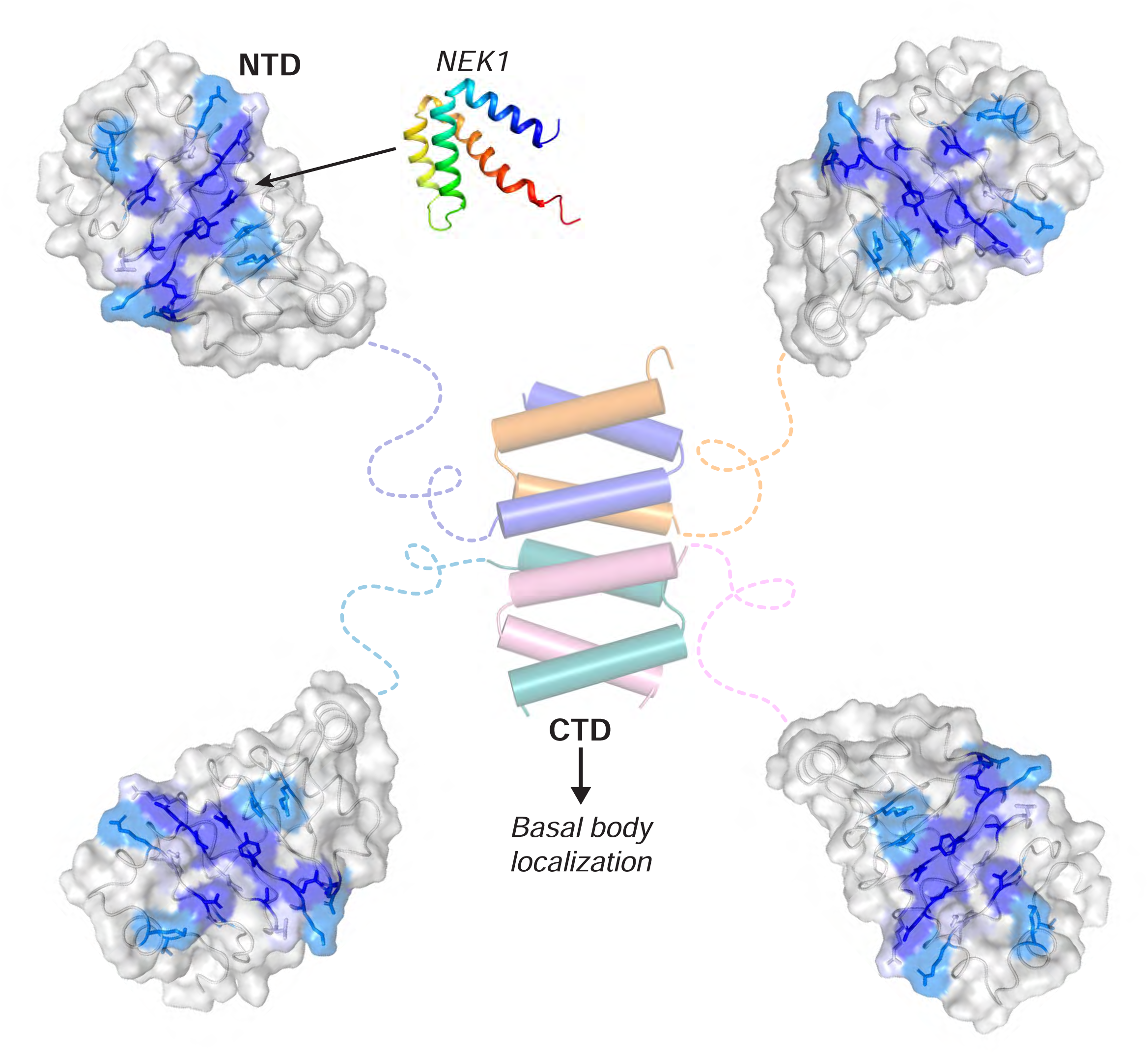
Bimodular organization of CFAP410. CFAP410-CTD forms a tetrameric helical bundle controlling the correct localization of the protein in the cell. CFAP410-NTD folds into a monomeric LRR motif and is loosely connected to the C-terminal bundle via a long disordered loop. The conserved surface patch (colored in blue) is likely involved in its interaction with NEK1 and other partners and thus plays an important role in the *in vivo* function of CFAP410.

## Materials and Methods

### Cloning and site-directed mutagenesis of TbCFAP410

Sequence encoding full-length TbCFAP410 was amplified from *T. brucei* genomic DNA using PCR and ligated into the pET15b vector (Novagen). All truncated constructs were subsequently generated from the full-length construct using either pET15b or a custom vector that provides a maltose-binding protein (MBP) and His10 tag to the N-terminus of the target proteins. The MBP-His10 tag can be cleaved off by the tobacco etch virus (TEV) protease. All single- and multi-residue mutations of TbCFAP410-NTD were generated using the megaprimer-based mutagenesis strategy as reported [22]. Incorporation of mutations was confirmed by DNA sequencing.

### Protein expression and purification

All recombinant TbCFAP410 proteins were expressed in *E. coli* (strain BL21-DE3). The transformed cells were grown in Luria-Bertani (LB) medium at 37°C to an OD600 of 0.6-0.8, and then subjected to cold shock on ice for 30 min. Protein expression was induced by addition of 0.5 mM isopropyl β-D-1-thiogalactopyranoside (IPTG), and cell cultures were further incubated at 18°C overnight. Cells were harvested by centrifugation in a Sorvall GS3 rotor (6,000×g, 12 min, 4°C). Cell pellets were resuspended in 10 ml of lysis buffer [20 mM HEPES pH 7.5, 100 mM NaCl, 20 mM imidazole, 1mM β-Mercaptoethanol, 1 mM phenylmethylsulfonyl fluoride (PMSF)] per L of cell culture.

Resuspended cells were lysed in an EmulsiFlex-C3 homogenizer (Avestin) at 40 bars and cell debris was pelleted by centrifugation (40,000×g, 40 min, 4°C). The supernatant was filtered (0.45-µm pore size, Amicon) and loaded onto a Ni-HiTrap® HP column pre-equilibrated with the same lysis buffer in an ÄKTA™ Pure chromatography system (Cytiva). The column with bound proteins was washed with 2× column volume (cv) of the same lysis buffer. Bound protein was subsequently eluted using a linear gradient concentration of imidazole (20 - 500 mM, 20×cv) in the lysis buffer.

The N-terminal MBP-His10 tag of all TbCFAP410 truncated proteins were cleaved off by incubating the pooled fractions of interest (20-40 ml) with approximately 1% (w/w) of the TEV protease in a dialysis bag against 2-4 L of dialysis buffer containing 20 mM HEPES (pH 7.5) and 100 mM NaCl (4°C, overnight). The dialyzed samples were loaded on a Ni-HiTrap® HP column to get rid of the cleaved tag and any uncut proteins. Target proteins in the flow-through were further purified on a Superdex-200 16/60 column (Cytiva) with a running buffer containing 20 mM HEPES (pH 7.5), 100 mM NaCl, and 1 mM dithiothreitol (DTT). The eluted proteins were then pooled and concentrated, and final concentration was determined using Ultraviolet (UV) absorbance at 280 nm. The samples were either used freshly or stored at -80°C for subsequent experiments.

### Crystallization and structural determination

Purified TbCFAP410-NTD proteins were subjected to extensive crystallization trials using several commercial crystallization kits from Hampton Research. Conditions of the initial hits were further optimized to obtain single crystals. The final condition for crystallizing TbCFAP410-NTD contained 0.1 M sodium citrate (pH 5.5) and 1 M NH4H2PO4 at 22 °C. X-ray diffraction data were collected at the ESRF synchrotron site and processed using XDS [23]. Crystal structure was determined by the molecular replacement method using a homology model of TbCFAP410 (aa16-138) generated by SWISS-MODEL [11] based on the crystal structure of the *C. reinhardtii* axonemal dynein light chain-1 (PDB code: 6L4P) [12]. The resulting model was optimized by multiple rounds of manual rebuilding in COOT [24] and refinement in Phenix [25], and finally validated in MolProbity [26]. Details of data collection and refinement are summarized in **Table 1**.

### Generation of constructs for *in vivo* assays in *T. brucei*

DNA sequence of TbCFAP410 was recoded in wild-type and all mutations for *in vivo* assays. The recoded DNA sequences were chemically synthesized (Twist Bioscience) and cloned into the plasmid pDEX577-mNG. All constructs were subsequently linearized with Not I and then electroporated using a BTX Gemini Twin Wave with 3 pulses of 1.7 kV for 100 µs as previously described [27]. Cells were recovered in SDM-79 for 8 hours before selection with phleomycin (5 µg/ml).

### Cell culture of *T. brucei*

SmOxP9 procyclic *T. brucei* expressing TbSAS6 endogenously tagged with mScarlet were used for all experiments [28]. These were derived from TREU 927 strain, expressing T7 RNA polymerase and tetracycline repressor [29] and were grown at 28°C in SDM-79 medium supplemented with 10% FCS [30]. Cell concentration was determined in a Z2 Coulter Counter particle counter.

### Fluorescence microscopy

Cells were induced overnight with doxycycline (0.5 µg/ml) and harvested by centrifugation at 800×g for 5 min. For live-cell microscopy, cells were washed two times in PBS supplemented with 10 mM glucose and 46 mM sucrose (vPBS). In the last wash, the DNA was stained using 10 µg/ml Hoechst 33342 (Sigma-Aldrich), re-suspended in vPBS and then mounted with a coverslip and immediately imaged. For cytoskeletons, cells were washed in PBS, settled onto glass slides for 5 min and treated with 1% (v/v) NP-40 in PEME for 1 min. Cytoskeletons were then fixed in −20°C methanol for 20 min and rehydrated in PBS for 10 min. After fixation, the DNA was stained with 20 μg/ml Hoechst 33342 in PBS, washed in PBS for 10 min or overnight and mounted before imaging. Images were taken using a Zeiss Axio Imager.Z1 microscope equipped with an ORCA-Flash 4.0 CMOS camera using a Plan-Apochromat 63×/1.4 NA oil objective. Images were acquired and analysed with ZEN 2 PRO software and assembled for publication in Adobe Illustrator CS6. For each induction 2 sets of 100 cytoskeletons were counted for each induced cell line using ImageJ (NIH, MD).

### Western blotting

After overnight induction with doxycycline (0.5 µg/ml), 2×10^7^ parasites were harvested by centrifugation at 800 × g for 3 min and washed twice with vPBS. Parasites were resuspended into 50 µl vPBS and 50 µl of 2 × Laemmli buffer, heated at 95 °C for five min and stored at -20 °C. Later, an amount equivalent to 4 × 10^6^ parasites was loaded onto a 10% (w/v) SDS-PAGE gel and electrophoresed before being transferred to a PVDF membrane. The membrane was blocked with 5% (w/v) milk in Tris-buffered saline with Tween-20 (TBS-tween) for one hour at room temperature, followed by overnight incubation with BB2 (anti-Ty) (1:100 in 5% milk) at 4 °C. After incubation, the membrane was washed five times for 5 min with TBS-tween, followed by incubation with the secondary antibody anti-mouse HRP (1:5000 in 5% milk) for one hour at room temperature. Finally, the membrane was washed five times, incubated with the Chemiluminescent Horseradish Peroxidase substrate for 2 min, and visualized using a G:BOX Chemi XRQ machine (Syngene).

### *C. reinhardtii* strain and culture growth

*C. reinhardtii* strain CC-5325 was grown photoheterotrophically in tris-acetate-phosphate (TAP) medium at 25°C under constant illumination at 50 µmol photons·m^-2^·s^-1^.

### *Sp*Cas9 protein purification and *In vitro* transcription of sgRNA

*Sp*Cas9 protein purification were performed as described [31]. The sgRNA target site (GTGAAGAACCTCAACCTCTGGGG) for the CrCFAP410 gene (Cre14.g625550) in *C. reinhardtii* was selected using CRISPR RGEN Tools (http://www.rgenome.net/cas-designer/). Transcription of sgRNA was accomplished using the HiScribe™ T7 Quick High Yield RNA Synthesis Kit (NEB) and Monarch® RNA Cleanup Kit (NEB).

### *In vitro* digestion of target DNA with RNP

Cells were picked into 50 μl 5% (w/v) Chelex100, heated to 95°C for 10 min and then on the ice for 1 min to generate crude genomic DNA. The template DNA contained the Cas9 cleavage site was amplified using primers (PAM-F: 5’-TGTCACTTTCACCTGCTGCGA-3’; PAM-R: 5’-CCAAGTTCGGCAGCTACAGT-3’) with the crude genomic DNA as a template. *In vitro* cleavage assays were according to the previous protocol with the following modifications [32]. 500 ng gRNA and 700 ng Cas9 were preincubated in Cas9 cleavage buffer (20 mM HEPES pH 7.5, 100 mM NaCl, 5 mM MgCl_2_, 0.5 mM DTT, 0.1 mM EDTA) at 37°C for 15 min, then 500 ng template DNA was added, and the mixtures were incubated at 37°C for 1 h, then 10 × STOP solution (0.5 M EDTA, 80% (v/v) glycerol, and 10% (w/v) SDS) was added. The final products were separated using 2% (w/v) agarose gel electrophoresis.

### Preparation of dsDNA donor containing hygromycin resistance cassette with microhomologies arms for targeting CrCFAP410

The hygromycin resistance cassette was amplified from the plasmid pHK266 [33]. The 20 bp microhomologies based on the flank sequence of the Cas9 cleavage site were added to both sides of the hygromycin resistance cassette by PCR using primers (donor-F: 5’-GAGGTGAAGAACCTCAACCTTCTTTCTTGCGCTATGACACT-3’; donor-R: 5’-CGTCCAGGTCCTGGCCCCAGTGGAGAAAGAGGCCAAAATC-3’). The PCR product was purified using Gel Extraction Kit (QIAGEN).

### Transformation of *C. reinhardtii* cells

Cells were cultured for 3 days to a cell density of 2 × 10^6^ cells/ml in TAP medium. The cells were then concentrated to 2 × 10^8^ cells/ml in TAP medium with 60 mM sorbitol. For each shot, Cas9 protein (50 µg, 0.53 nmol) and sgRNA (1.6 nmol) as 1:3 molar ratio at 37°C for 15 min to assemble RNP complexes. 20 µl RNP and 2 µg dsDNA donor were added to 80 µl cell cultures in the 4 mm cuvette. Electroporation was conducted using BTX ECM® 630 Electroporation System (600 V, 50 µF, 350Ω). Immediately after electroporation, 600 µl of TAP medium with 60 mM sorbitol was added. Cells were recovered overnight (24 h) in 10 ml TAP with 60 mM sorbitol and plated onto TAP media supplemented with 20 µg/ml hygromycin. Colonies appeared after 5–7 days.

### Genotyping and sequencing of potential mutants

To determine whether the dsDNA donor inserted into the cleavage site, the crude genomic DNA from colonies was used as templates, and PAM-F and PAM-R were used as PCR primers. The PCR products were then analyzed using 1% (w/v) agarose gel. For sequencing of CrCFAP410 mutations, about 2,188 bp PCR bands were gel purified and sequenced using the primers PAM-F and PAM-R.

### Flagellar percentage and length measurements

The methods of flagellar length measurements and percentage of the flagellated cells were described previously [34]. For each sample, at least 100 cells and 200 flagella were measured.

### Molecular dynamics (MD) simulations

Atomistic models for wild-type TbCFAP410-NTD and all mutants (V37F, C63Y, R75P, Y107C, Y107H and V111M) were constructed starting from the crystal structure by *in silico* mutagenesis using Pymol (http://www.pymol.org). All further calculations were performed using GROMACS 2019 package [35] and Amber99SB-ILDN force-field parameters [36]. In all cases, the simulated system was placed into a 7×7×7-nm^3^ octahedral box, energy minimized and solvated with TIP3P water molecules. Additionally, Na^+^ and Cl^−^ ions were added to achieve electroneutrality at the final salt concentration of 150 mM. The complete system was then energy-minimized using position restraints placed on protein Cα-atoms, and subjected to 5,000 steps of NVT molecular dynamics (MD) with a 0.5-fs time step and subsequent 250,000 steps of NPT MD with a 1-fs time step. After this initial equilibration, a 1 μs NPT production MD run was simulated (5×10^8^ steps with a 2-fs time step) for each system. All analysis was performed on the final 750 ns of each simulated trajectory. A twin-range (10 Å, 10.5 Å) spherical cut-off function was used to truncate the van der Waals interactions. Electrostatic interactions were treated using the particle-mesh Ewald summation (real space cut-off of 10-Å and 1.2-Å grid with fourth-order spline interpolation). MD simulations were carried out using three-dimensional periodic boundary conditions, in the isothermal−isobaric (NPT) ensemble with an isotropic pressure of 1.013 bar and a constant temperature of 295 K. The pressure and temperature were controlled using a Nose-Hoover thermostat [37, 38] and a Parrinello–Rahman barostat with 0.5-ps and 10-ps relaxation parameters, respectively, and a compressibility of 4.5×10^−5^ bar^−1^ for the barostat.

### Accession code

Coordinates and structure factors have been deposited in the Protein Data Bank (PDB) under accession code 8AXJ.

## Acknowledgements

The authors are grateful to the staff at the beamlines of ID23-1 at the European Synchrotron Radiation Facility (ESRF) for their help with X-ray diffraction data collection.

This work was supported by the grant I5960-B2 from the Austrian Science Fund (FWF) to GD and a Newton International Fellowship (NIF\R1\191618) to HG. SA acknowledges the VIP2 fellowship 2020 funded by the European Union’s Horizon 2020 research and innovation programme under the Marie Skłodowska-Curie grant agreement No. 847548. JDS acknowledges the Wellcome Trust and Oxford Brookes University for their support.

