## Supplemental information for "CFAP410 has a bimodular architecture with a conserved surface patch on its N-terminal LRR motif for binding interaction partners"

#### Legends for supplemental figures

**Figure S1. Structural analysis of the interaction between the extra loop and the LRR motif.** (**A**) 2D plot of the interaction between the C-terminal loop (aa139-160) and the LRR motif of TbCFAP410-NTD. The plot was generated using DIMPLOT in the LigPlot plus suite. Residues involved in hydrogen bond formation are shown as ball-and-sticks, with oxygen, nitrogen and carbon atoms colored in red, blue, and gray, respectively. Dotted lines in cyan indicate hydrogen bonds. Non-bonded residues involved in hydrophobic interactions are shown as spoked arcs. (**B**) Sequence alignment of TbCFAP410 and homologs in their NTDs. Residues involved in hydrogen bond hydrophobic interactions as shown in (A) are indicated by arrows and dots, respectively. (**C**) Two orthogonal views of the crystal structure of TbCFAP410-NTD, with the LRR motif (aa1-138) shown as semitransparent surface and the loop as green cartoon and sticks. Highly and relatively conserved residues on the LRR motif are shaded in blue and light cyan, respectively. (**D**) Open-book views of the interfaces on the LRR and the loop. $2F_o$-$F_c$ electron density maps are shown (purple) for the entire loop at 2 σ level.

**Figure S2. Conservation analysis of the NTDs of various CFAP410 homologs.** (**A**) Sequence alignment of homologs of *H. sapiens*, *T. brucei* and *C. reinhardtii* CFAP410 proteins. Highly conserved surface residues are indicated by arrows under the alignment (dark, ocean and light blue are for residues with >90%, 80-90% and 70-80% identities, respectively). Residues corresponding to the disease-causing mutations are marked by diamonds with the residue numbers corresponding to their position in the TbCFAP410 sequence. (**B** & **C**) Mapping the conserved residues onto the crystal structure of

TbCFAP410-NTD. The solvent-exposed flat surface on the β sheet contains many conserved residues (B), whereas the opposite side of the LRR motif with the extra loop is devoid of conserved residues (C).

**Figure S3. Homology modeling of HsCFAP410-NTD and its comparison with TbCFAP410-NTD.** (**A**) NCBI-BLAST report for the NTD of *H. sapiens* and *T. brucei* CFAP410. Residues with mutations that cause diseases in human are shaded in cyan. (**B**) Ribbon diagram of the crystal structure of TbCFAP410-NTD (blue) superimposed with the homology models of HsCFAP410-NTD generated by either SWISS-MODEL (green) or AlphaFold2 (pink). The five residues highlighted in (A) are shown as sticks and labeled. (**C**) SWISS-MODEL report for the HsCFAP410-NTD homology modeling. (**D**) Predicted alignment error (PAE) plots of AlphaFold2 homology models of HsCFAP410-NTD. (**E**) Predicted local distance difference test (pLDDT) plots of the AlphaFold2 homology models of HsCFAP410-NTD. It is a per-residue measure of local confidence on a scale from 0-100 (the higher the better).

**Figure S4. Simulation and prediction analyses of wild-type and mutant TbCFAP410-NTD.** (**A**) Conformational analysis of the C63Y mutant simulations. *Left:* central configuration of the most populated cluster in the hNTD C63Y simulations (cluster occupancy 21%); *Right:* central configuration of the second most populated cluster in the C63Y simulations showing the large opening of the C-terminal cleft, which coincides with the formation of the E61-K156 salt bridge. (**B**) Analysis of hNTD wild-type (WT) and mutant simulations with the number of conformational clusters with cluster occupancy P >

1%; cluster occupancy of the most populated cluster ($P_{MPC}$); all-atom RMSD between the central structures of the most populated clusters in mutant and wildtype simulations; change in conformational entropy upon mutation ($DS_{conf} = S_{mutant}-S_{wt}$) whereby $S_{mutant}$ was rescaled by the relative number of degrees of freedom; change in the total solvent-accessible surface area ($DSASA_T$) and its hydrophilic ($DSASA_{Hphi}$) and hydrophobic components ($DSASA_{Hpho}$) upon mutation ($DSASA = SASA_{mutant}-SASA_{wt}$). (**C**) Superposition of TbCFAP410-NTD crystal structure with five models of the mutant C63Y generated by AlphaFold. The bulky aromatic sidechain of tyrosine in the mutant causes the flip of the helix outwards (insert) that subsequently destabilizes the interaction with the loop.

**Figure S5. Comparison of TbCFAP410 with its paralog.** (**A**) Sequence alignment of TbCFAP410 and its paralog from *T. brucei*. Residues in the conserved surface patch on TbCFAP410-NTD are indicated by arrows (green, conserved in paralog; orange, different in paralog). (**B**) Superposition of the AlphaFold model of the paralog on the crystal structure of TbCFAP410-NTD. (**C**) The conserved surface patch on TbCFAP410-NTD. (**D**) Conservation analysis of the surface patch on TbCFAP410-NTD versus its paralog.

**Figure S6. Sequence comparison of all HsCFAP410 isoforms.** (**A**) Sequence alignment of all 11 isoforms (i.e. 1-4 and X1-X7) of HsCFAP410. The ranges of both NTD and CTD are indicated. The alignment was carried out using the option of "Muscle with defaults" in Jalview. Residues are shaded by the degree of conservation. (**B**) Two orthogonal views of the three-dimensional structure of a tetrameric model of HsCFAP410 generated by AlphaFold. The N- and C-terminal regions are colored orange and teal, respectively.

Figure S1

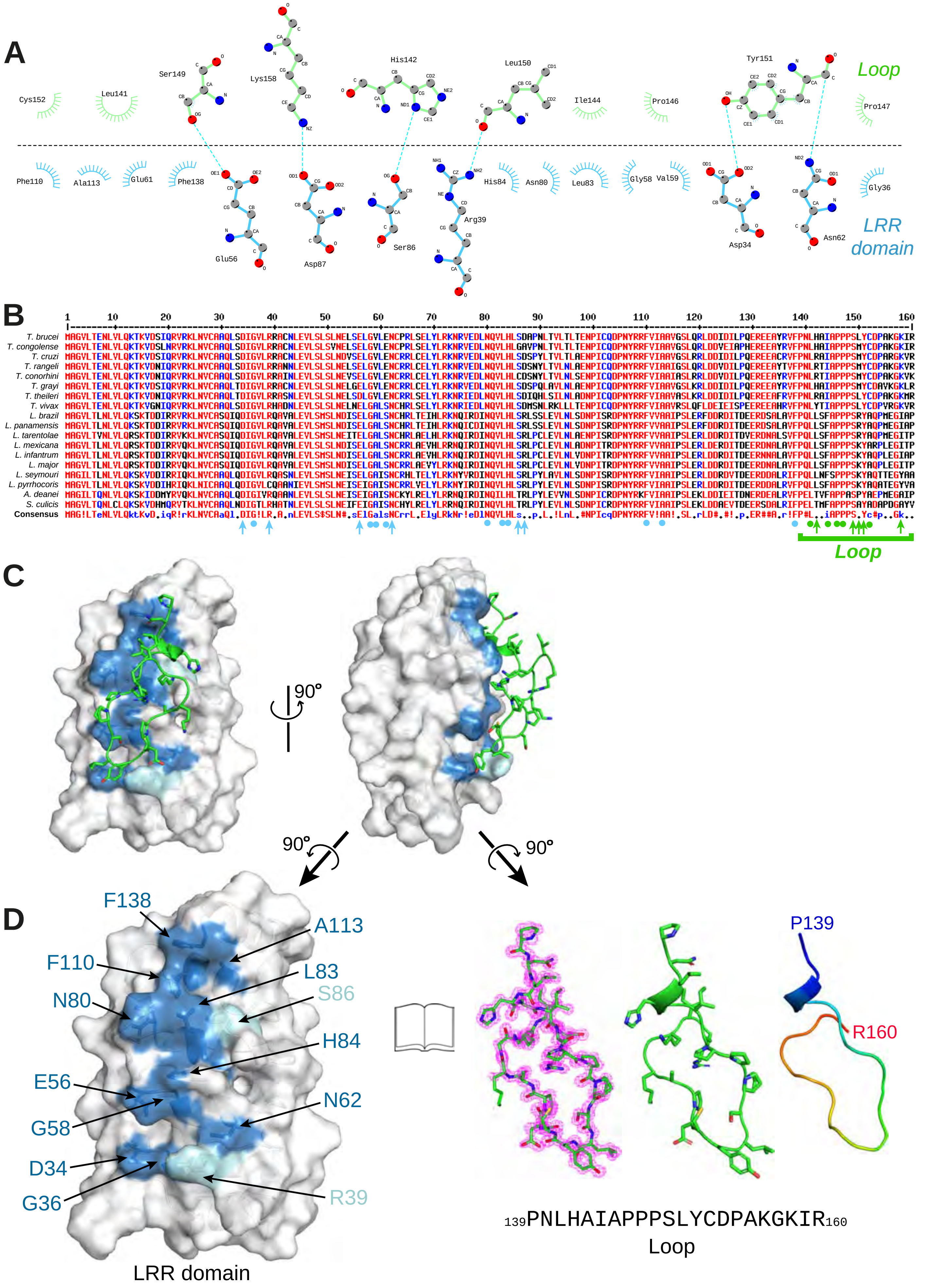

A
Loop
LRR domain
Cys152
Leu141
Ser149
Lys158
His142
Leu150
Ile144
Pro146
Tyr151
Pro147
Phe110
Ala113
Glu61
Phe138
Glu56
Asp87
Ser86
Arg39
His84
Asn80
Leu83
Gly58
Val59
Asp34
Asn62
Gly36
B
T. brucei
T. congolense
T. cruzi
T. rangeli
T. conorhini
T. grayi
T. theileri
T. vivax
L. brazil
L. panamensis
L. tarentolae
L. mexicana
L. infantrum
L. major
L. seymouri
L. pyrrhocoris
A. deanei
S. culicis
Consensus
Loop
C
90°
D
F138
A113
F110
L83
N80
S86
H84
E56
N62
G58
D34
G36
R39
P139
R160
LRR domain
139PNLHAIAPPPSLYCDPAKGKIR160
Loop

Figure S2

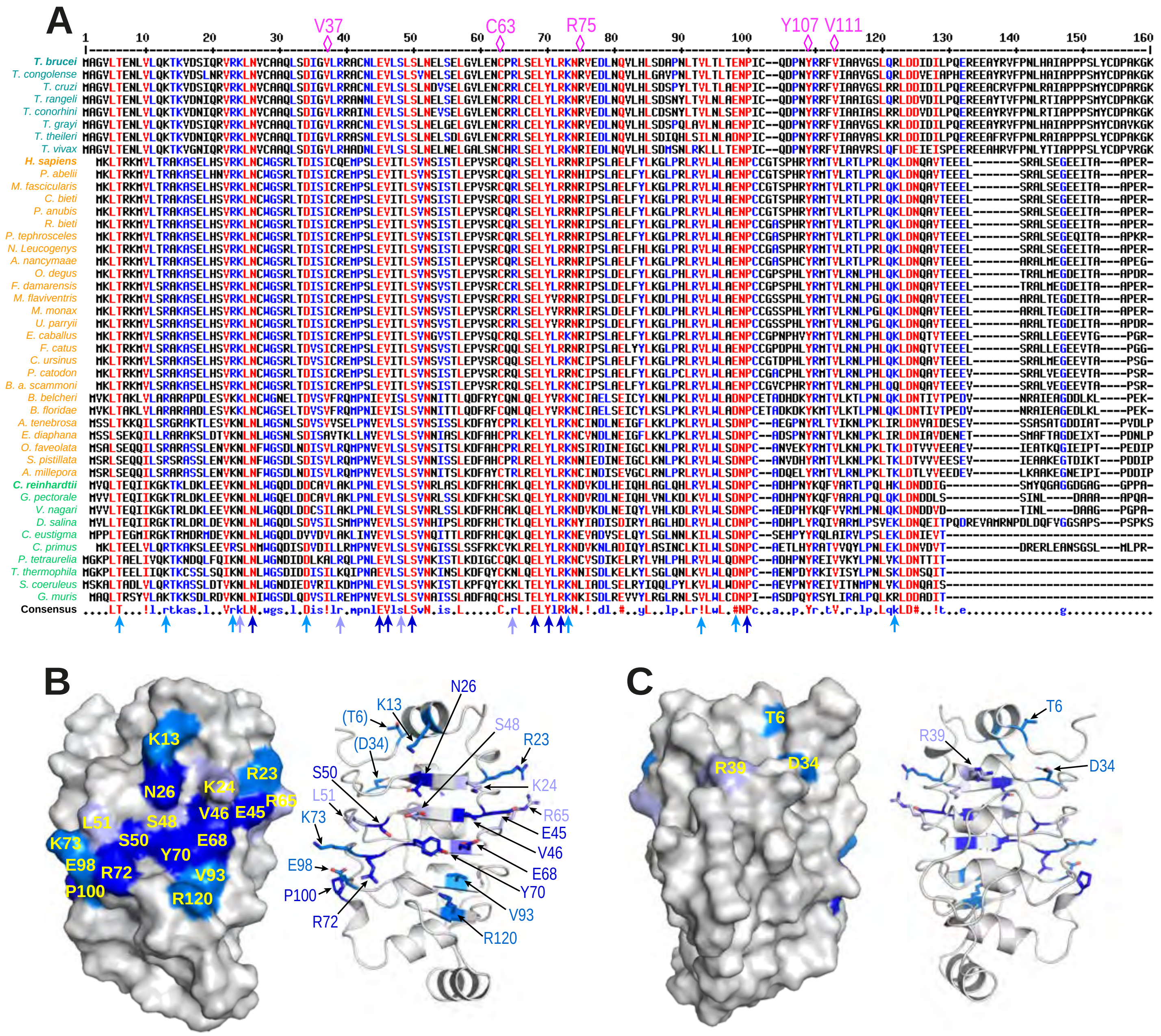

A
V37
C63
R75
Y107
V111
T. brucei
T. congolense
T. cruzi
T. rangeli
T. conorhini
T. grayi
T. theileri
T. vivax
H. sapiens
P. abelii
M. fascicularis
C. bieti
P. anubis
R. bieti
P. tephrosceles
N. Leucogenys
A. nancymaae
O. degus
F. damarensis
M. flaviventris
M. monax
U. parryii
E. caballus
F. catus
C. ursinus
P. catodon
B. a. scammoni
B. belcheri
B. floridae
A. tenebrosa
E. diaphana
O. faveolata
S. pistillata
A. millepora
C. reinhardtii
G. pectorale
V. nagari
D. salina
C. eustigma
C. primus
P. tetraurelia
T. thermophila
S. coeruleus
G. muris
Consensus
B
K13
N26
K24
R23
L51
S48
V46
E45
R65
K73
S50
Y70
E68
E98
R72
V93
P100
R120
(T6)
(D34)
S48
R23
S50
K24
L51
R65
K73
E45
V46
E98
E68
P100
Y70
R72
V93
R120
C
T6
R39
D34
T6
R39
D34

### Figure S3

**A**

| Score | Expect | Method | Identities | Positives | Gaps |
|---|---|---|---|---|---|
| 93.2 bits (230) | 5e-23 | Compositional matrix adjust. | 52/128 (**41%**) | 78/128 (60%) | 2/128 (1%) |

```
T.brucei    5  LTENLVLQKTKVDSIQRVRKLNVCAAQLSDIGVLRRACNLEVLSLSLNELSELGVLENCPRLSE   68
               LT  +VL + K    +   VRKLN    ++L+DI + +    +LEV++LS+N +S L   +   C
H.sapiens   3  LTRKMVLTRAKASELHSVRKLNCWGSRLTDISICQEMPSLEVITLSVNSISTLEPVSRCQRLSE   66

T.brucei   69  LYLRKNRVEDLNQVLHLSDAPNLTVLTLTENPIC-QDPN-YRRFVIAAVGSLQRLDDIDILPQE   130
               LYLR+NR+   L ++ +L    P L VL L ENP C    P+ YR  V+   +  LQ+LD+   +   +E
H.sapiens  67  LYLRRNRIPSLAELFYLKGLPRLRVLWLAENPCCGTSPHRYRMTVLRTLPRLQKLDNQAVTEEE   130
```

**B**

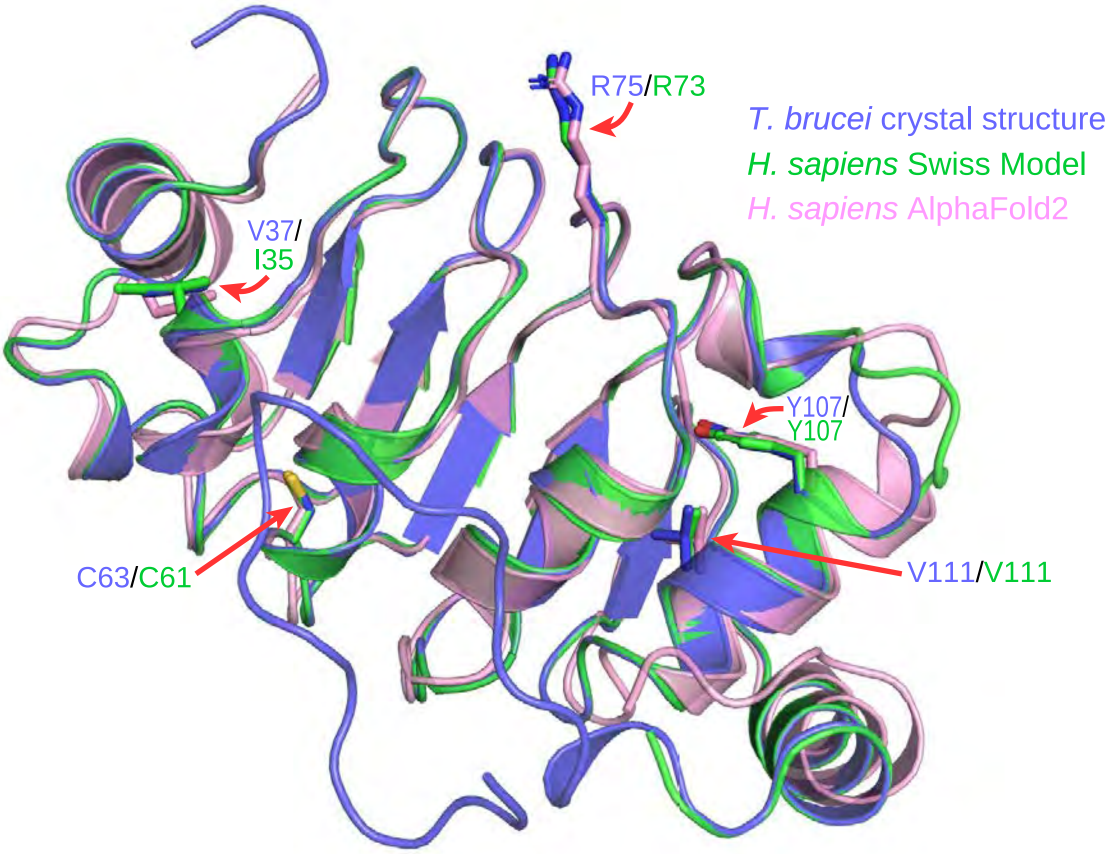


**C** **D** **E**

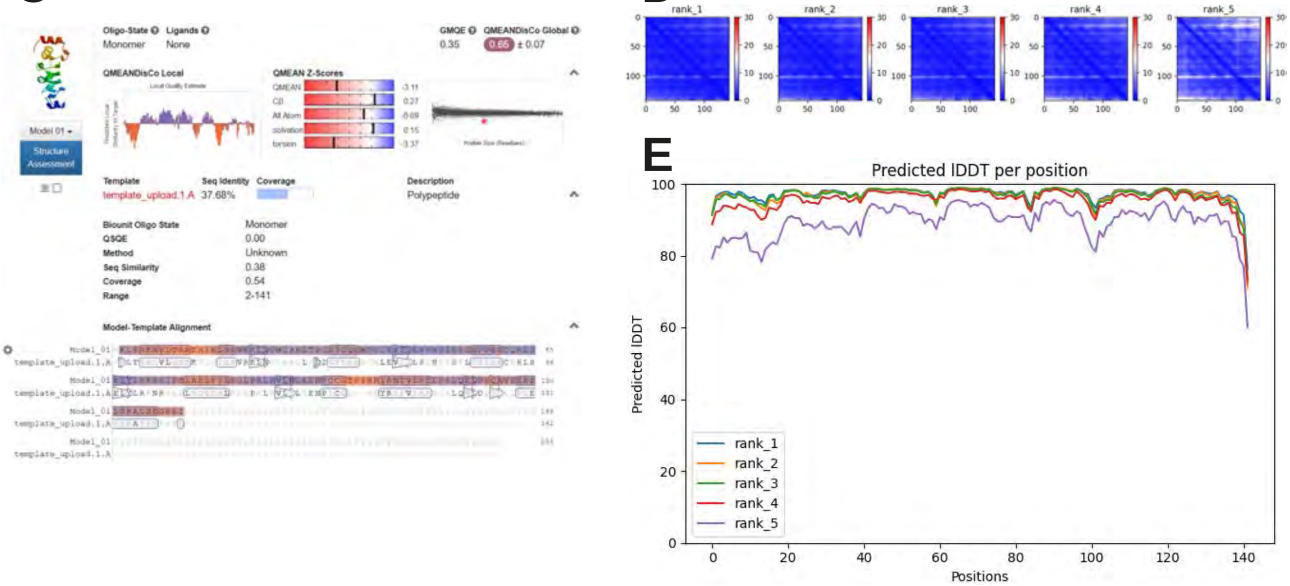

### Figure S4

**A**

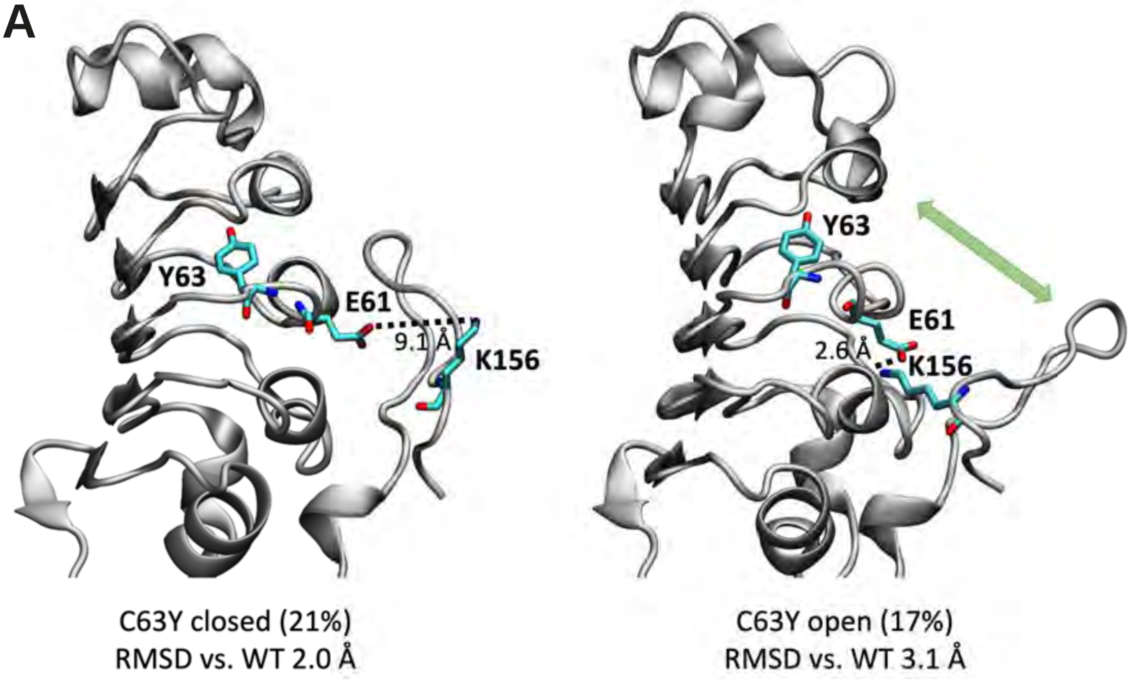


**B**

| system | # clusters (P > 1%) | $P_{MPC}$ | RMSD | $\Delta S_{conf}$ | $\Delta SASA_T$ | $\Delta SASA_{Hphi}$ | $\Delta SASA_{Hpho}$ |
|---|---|---|---|---|---|---|---|
| WT | 6 | 59.1 | - | - | - | - | - |
| V37F | 1 | 99.9 | 0.9 | -34.1 | -1.0 | 0.0 | -1.0 |
| C63Y | 10 | 20.5 | 2.0 | 15.4 | 2.0 | 0.5 | 1.5 |
| R75P | 5 | 59.2 | 1.1 | 8.1 | -1.0 | -1.5 | 0.5 |
| Y107C | 8 | 31.3 | 1.1 | 25.7 | 0.5 | 0.3 | 0.2 |
| Y107H | 7 | 76.0 | 1.1 | 3.2 | 0.5 | 0.5 | 0.0 |
| V111M | 2 | 94.3 | 0.9 | 2.9 | 0.0 | 0.0 | 0.0 |
| | | % | Å | J/mol K | $nm^2$ | | |

**C**

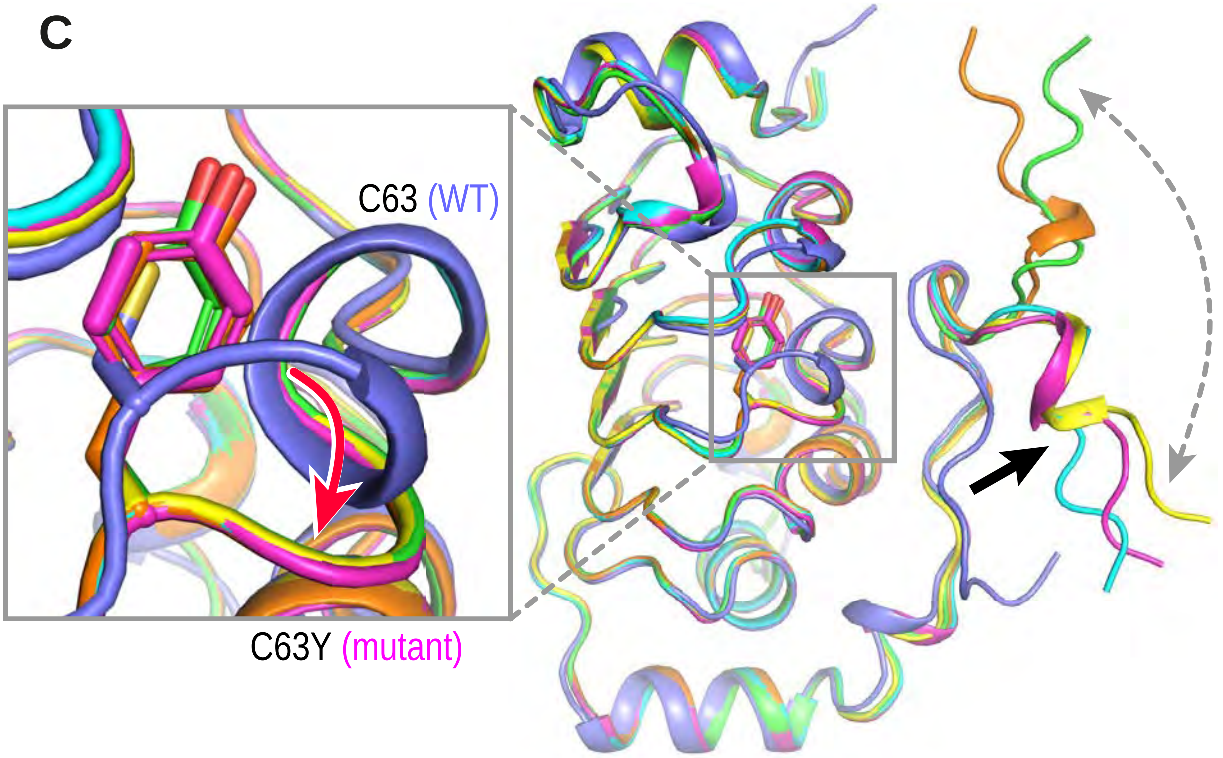

### Figure S5

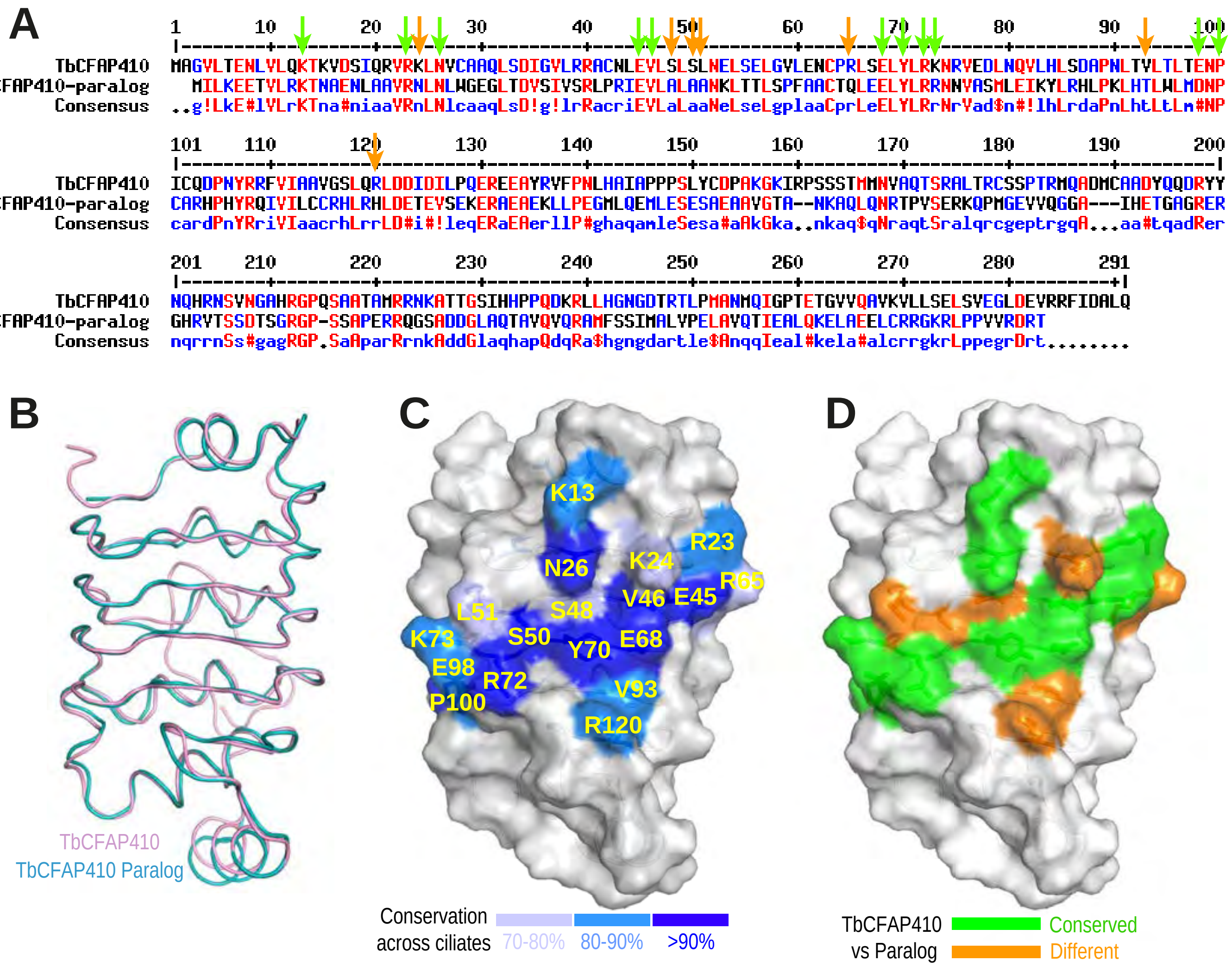

A
1 10 20 30 40 50 60 70 80 90 100
TbCFAP410 MAGVLTENLVLQKTKVDSIQRVRKLNVCAAQLSDIGVLRRACNLEVLSLSLNELSELGVLENCPRLSELYLRKNRVEDLNQVLHLSDAPNLTVLTLTENP
TbCFAP410-paralog MILKEETVLRKTNAENLAAVRNLNLWGEGLTDVSIVSRLPRIEVLALAANKLTTLSPFAACTQLEELYLRRNNVASMLEIKYLRHLPKLHTLWLMDNP
Consensus ..g!LkE#lVLrKTna#niaaVRnLNlcaaqLsD!g!lrRacriEVLaLaaNeLseLgplaaCprLeELYLRrNrVad$n#!lhLrdaPnLhtLtLm#NP
101 110 120 130 140 150 160 170 180 190 200
TbCFAP410 ICQDPNYRRFVIAAVGSLQRLDDIDILPQEREEAYRVFPNLHAIAPPPSLYCDPAKGKIRPSSSTMMNVAQTSRALTRCSSPTRMQADMCAADYQQDRYY
TbCFAP410-paralog CARHPHYRQIVILCCRHLRHLDETEVSEKERAEAEKLLPEGMLQEMLESESAEAAVGTA--NKAQLQNRTPVSERKQPMGEVVQGGA---IHETGAGRER
Consensus cardPnYRriVIaacrhLrrLD#i#!leqERaEAerllP#ghaqamleSesa#aAkGka..nkaq$qNraqtSralqrcgeptrgqA...aa#tqadRer
201 210 220 230 240 250 260 270 280 291
TbCFAP410 NQHRNSVNGAHRGPQSAATAMRRNKATTGSIHHPPQDKRLLHGNGDTRTLPMANMQIGPTETGVVQAVKVLLSELSVEGLDEVRRFIDALQ
TbCFAP410-paralog GHRVTSSDTSGRGP-SSAPERRQGSADDGLAQTAVQVQRAMFSSIMALVPELAVQTIEALQKELAEELCRRGKRLPPVVRDRT
Consensus nqrrnSs#gagRGP.SaAparRrnkAddGlaqhapQdqRa$hgngdartle$AnqqIeal#kela#alcrrgkrLppegrDrt........
B
TbCFAP410
TbCFAP410 Paralog
C
K13
R23
N26
K24
R65
V46
E45
L51
S48
K73
S50
Y70
E68
E98
R72
V93
P100
R120
Conservation across ciliates
70-80%
80-90%
>90%
D
TbCFAP410 vs Paralog
Conserved
Different

### Figure S6

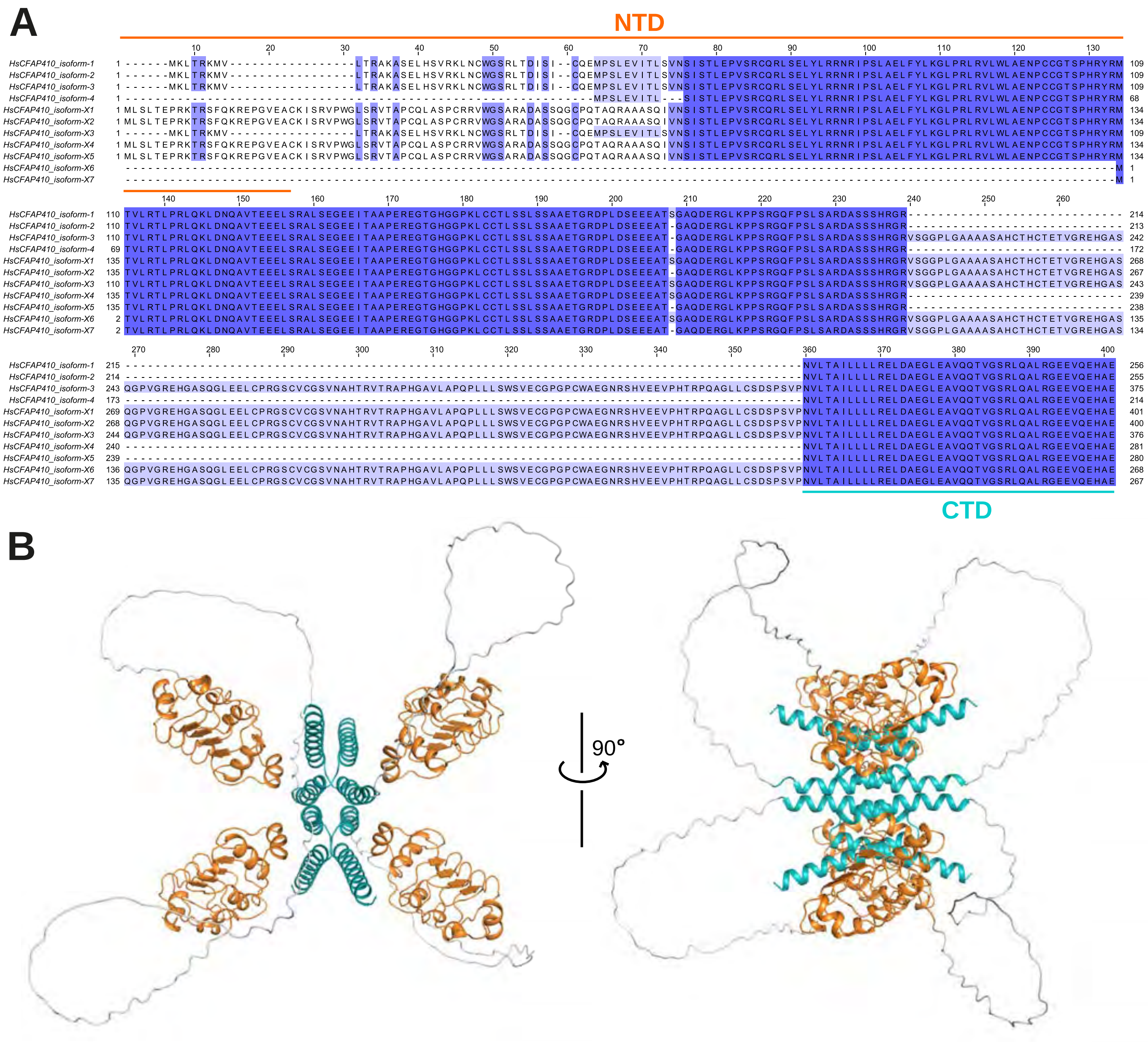
